# Deep evolutionary analysis reveals the design principles of fold A glycosyltransferases

**DOI:** 10.1101/2019.12.31.891697

**Authors:** Rahil Taujale, Aarya Venkat, Liang-Chin Huang, Wayland Yeung, Khaled Rasheed, Arthur S. Edison, Kelley W. Moremen, Natarajan Kannan

**Affiliations:** Institute of Bioinformatics, University of Georgia, Athens, GA 30602; Complex Carbohydrate Research Center, University of Georgia, Athens, GA 30602; Departments of Biochemistry and Molecular Biology, University of Georgia, Athens, GA 30602; Departments of Computer Science, University of Georgia, Athens, GA 30602

**Keywords:** glycosyltransferases, GT evolution, common core, GT machine learning, donor prediction

## Abstract

Glycosyltransferases (GTs) are prevalent across the tree of life and regulate nearly all aspects of cellular functions by catalyzing synthesis of glycosidic linkages between diverse donor and acceptor substrates. Despite the availability of GT sequences from diverse organisms, the evolutionary basis for their complex and diverse modes of catalytic and regulatory functions remain enigmatic. Here, based on deep mining of over half a million GT-A fold sequences from diverse organisms, we define a minimal core component shared among functionally diverse enzymes. We find that variations in the common core and the emergence of hypervariable loops extending from the core contributed to the evolution of catalytic and functional diversity. We provide a phylogenetic framework relating diverse GT-A fold families for the first time and show that inverting and retaining mechanisms emerged multiple times independently during the course of evolution. We identify conserved modes of donor and acceptor recognition in evolutionarily divergent families and pinpoint the sequence and structural features for functional specialization. Using the evolutionary information encoded in primary sequences, we trained a machine learning classifier to predict donor specificity with nearly 88% accuracy and deployed it for the annotation of understudied GTs in five model organisms. Our studies provide an evolutionary framework for investigating the complex relationships connecting GT-A fold sequence, structure, function and regulation.

## Introduction

Complex carbohydrates make up a large bulk of the biomass of any living cell and play essential roles in biological processes ranging from cellular interactions, pathogenesis, immunity, quality control of protein folding and structural stability (1). Biosynthesis of complex carbohydrates in most organisms is carried out by a large and diverse family of Glycosyltransferases (GTs) that transfer sugars from activated donors such as nucleotide diphosphate and monophosphate sugars or lipid linked sugars to a wide range of acceptors that include saccharides, lipids, nucleic acids and metabolites. Nearly 1% of protein coding genes in the human genome, and more than 2% of the *Arabidopsis* genome, are estimated to be GTs. GTs have undergone extensive variation in primary sequence and three-dimensional structure to catalyze glycosidic linkages between diverse donor and acceptor substrates. However, an incomplete understanding of the relationships connecting sequence, structure, function and regulation presents a major bottleneck in understanding pathogenicity, metabolic and neurodegenerative diseases associated with abnormal GT functions (2, 3).

Structurally, GTs adopt one of three folds (GT-A, -B or -C) with the GT-A Rossmann like fold being the most common. The GT-A fold is characterized by alternating β-sheets and α-helices (α/β/α sandwich) found in most nucleotide binding proteins (4). Majority of GT-A fold enzymes are metal dependent and conserve a DxD motif in the active site that helps coordinate the metal ion and the nucleotide sugar. Currently, 109 GT-A families have been catalogued in the Carbohydrates Active Enzymes (CAZy) database (5). These families can be broadly classified into two categories based on their mechanism of action and the anomeric configuration of the glycosidic product relative to the sugar donor, namely, inverting or retaining. Inverting GTs generally employ an SN2 single displacement reaction mechanism that results in inversion of anomeric configuration for the product, whereas retaining GTs generally employ a dissociative SNi-type mechanism that retains the anomeric configuration of the product (6). While the sequence basis for inverting and retaining mechanisms is not well understood, most inverting GT-As have a conserved Asp or Glu within a xED motif that serves as the catalytic base to deprotonate the incoming nucleophile of the acceptor, and initiate nucleophilic attack with direct displacement of the phosphate leaving group (7, 8). Retaining GT-As bind the sugar donor similarly to the inverting enzymes, but shift the position of the acceptor nucleophile to attack the anomeric carbon from an obtuse angle using a phosphate oxygen of the sugar donor as the catalytic base and employ a dissociative mechanism that retains the anomeric linkage for the resulting glycosidic product (6). Such mechanistic diversity of GTs is further illustrated by recent crystal structures of GTs bound to acceptor and donor substrates which show that different acceptors are accommodated in the active site through variable loop regions emanating from the catalytic core (6). However, whether these observations hold for the entire super-family is not known because of the lack of structural information for the vast number of GTs.

The wealth of sequence data available on GTs provides an opportunity to infer underlying mechanisms through deep mining of large sequence datasets. In this regard, the CAZY database serves as a valuable resource for generating new functional hypotheses by classifying GT enzymes into individual families based on overall sequence similarity. However, a broader understanding of how these enzymes evolved to recognize diverse donor and acceptor substrates requires a global comparison of diverse GT-A fold enzymes. Such comparisons are currently a challenge due to limited sequence similarity between families and the lack of a phylogenetic framework to relate evolutionarily divergent families. Previous efforts to investigate GT evolution have largely focused on individual families or pathways (9, 10) and have not explicitly addressed the challenge of mapping the evolution of functional diversity across families.

Here through deep mining of over half a million GT-A fold related sequences from diverse organisms, and application of specialized computational tools developed for the study of large gene families (11, 12), we define a common core shared among diverse GT-A fold enzymes. Using the common core features, we generate a phylogenetic framework for relating functionally diverse enzymes and show that inverting and retaining mechanisms emerged independently multiple times during evolution. We identify convergent modes of substrate recognition in evolutionarily divergent families and pinpoint sequence and structural features associated with functional specialization. Finally, based on the evolutionary and structural features gleaned from a broad analysis of diverse GT-A fold enzymes, we develop a machine learning (ML) framework for predicting donor specificity with nearly 88% accuracy. We predict donor specificity for uncharacterized GT-A enzymes in diverse model organisms and provide testable hypotheses for investigating the relationships connecting GT-A fold structure, function and evolution.

## Results

### An ancient common core shared among diverse GT-A fold enzymes

To define common features shared among diverse GT-A fold enzymes, we generated a multiple sequence alignment of over 600,000 GT-A fold related sequences in the non-redundant (NR) sequence database (13) using curated multiple-aligned profiles of diverse GTs (Table S1). The alignment profiles were curated using available crystal structures (Methods)(14). The resulting alignment revealed a GT-A common core consisting of 231 aligned positions. These aligned positions are referred to throughout this analysis and are mapped to representative structures in Dataset S1. The common core is defined by eight β sheets and six α helices, including three β sheets and α helices from the N-terminal Rossmann fold (Fig. 1A,B).

**Figure 1:**
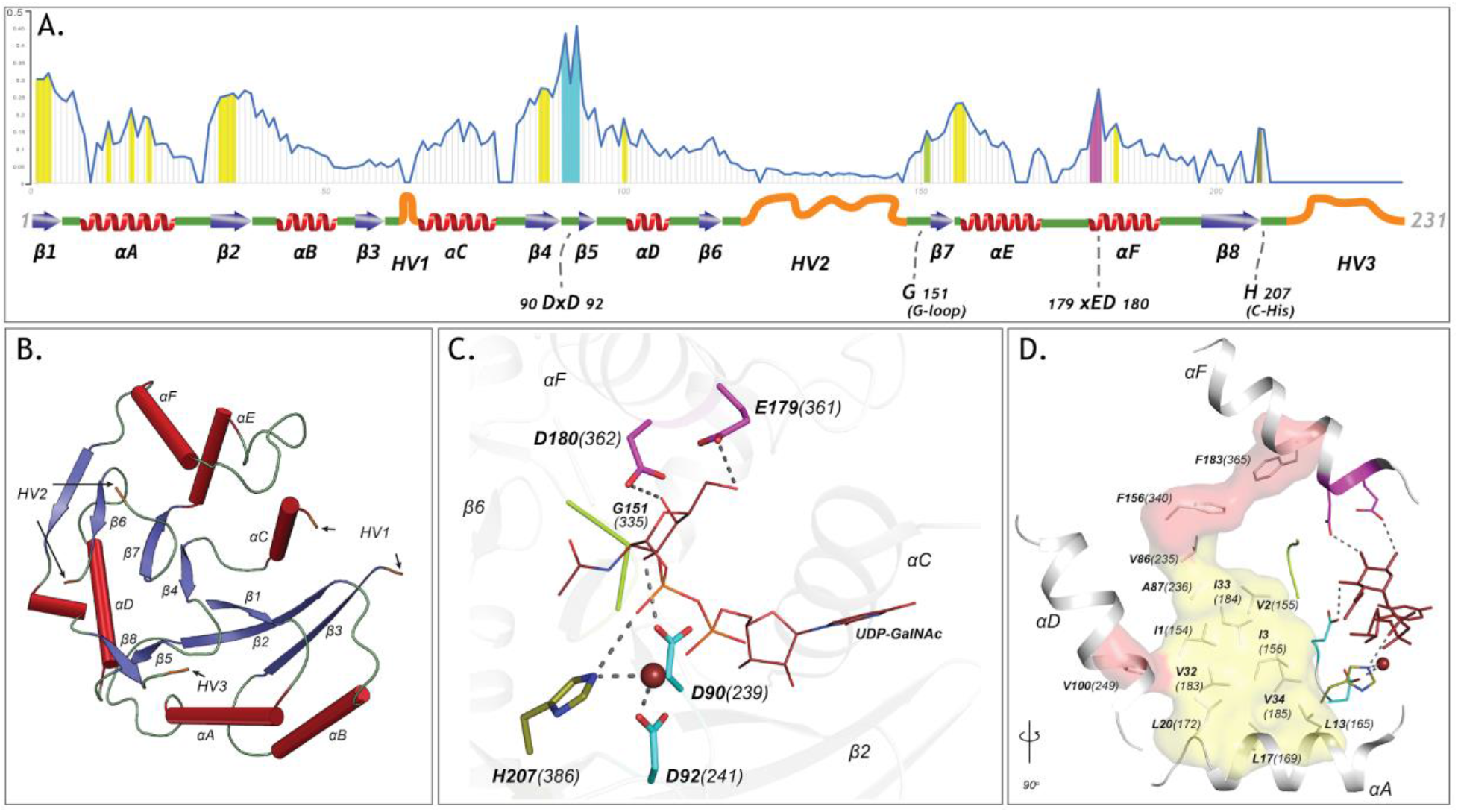
The GT-A common core and its elements. A) Plot showing the schematics of the GT-A common core with 231 aligned positions. Conserved secondary structures (red α-helices, blue β-sheets, green loops) and hypervariable regions (HVs)(orange) are shown. Conservation score for each aligned position is plotted in the line graph above the schematics. Evolutionarily constrained regions in the core: the hydrophobic positions (yellow) and the active site residues (DxD: Cyan, xED: Magenta, G-loop: green, C-His: olive) are highlighted above the positions. B) The conserved secondary structures and the location of HVs are shown in the N-terminal GT2 domain of the multidomain chondroitin polymerase structure from E. coli (PDB: 2z87) that is used as a prototype as it displays closest similarity to the common core consensus (SI Methods). C) Active site residues of the prototypic GT-A structure. Metal ion and donor substrate are shown as a brown sphere and sticks, respectively. D) Architecture of the hydrophobic core (Yellow: core conserved in all Rossmann fold containing enzymes, Red: core elements present only in the GT-A fold). Residues are labeled based on their aligned positions. Numbers within parentheses indicate their position in the prototypic (PDB: 2z87) structure.

Quantification of the evolutionary constraints imposed on the common core reveal twenty residues shared among diverse GT-A fold families. These include the DxD and the xED motif residues involved in catalytic functions, and other residues not typically associated with catalysis (Fig. 1A) such as the conserved glycine at aligned position 151 (G335 in 2z87) in the flexible G-loop and a histidine residue (H386 in 2z87) in the C-terminal tail at aligned position 207, henceforth referred to as the C-His. Residues from the G-loop in some families, such as the blood ABOs (GT6) and glucosyl-3-phosphoglycerate synthases (GpgS; GT81), contribute to donor binding (15, 16). The C-His, likewise, coordinates with the metal ion and contributes to catalysis in a subset of GTs, such as polypeptide N-acetylgalactosaminyl transferases (ppGalNAcTs; GT27) and lipopolysaccharyl-α-1,4-galactosyltransferase C (LgtC; GT8) (17, 18). The conservation of these residues across diverse GT-A fold enzymes suggest that they likely perform similar functional roles in other families as well.

The remaining core conserved residues include fourteen hydrophobic residues that are dispersed in sequence, but spatially cluster to connect the catalytic and donor binding sites in the Rossmann fold. Eleven out of the fourteen residues (highlighted in yellow in Fig. 1D) are shared by other Rossmann fold proteins (Fig. S1) suggesting a role for these residues in maintaining the overall fold. Three hydrophobic residues (V249, F340, F365; shown in red surface in Fig. 1D), however, are unique to GT-A fold enzymes, and structurally bridge the αF helix (containing the xED motif), the αD helix and the Rossmann fold domain. Although the functional significance of this hydrophobic coupling is not evident from crystal structures, in some families (GT15 and GT55) the hydrophobic coupling between αF and the Rossmann fold domain is replaced by charged interactions (Fig. S2). The structural and functional significance of these family specific variations are discussed below.

Our broad evolutionary analysis also reveals three hypervariable regions (HVs) extending from the common core. These include an extended loop segment connecting β3 strand and αC helix (HV1), a segment longer than 28 amino acids connecting β7 and β8 strand (HV2) and a C-terminal tail extending from the β8 strand (HV3) in the common core. These HVs, while conserved within families, display significant conformational and sequence variability across families (Fig. 1A, Fig. S3) and encode family-specific motifs that contribute to acceptor specificity in individual families, as discussed below.

### A phylogenetic framework relating diverse GT-A fold families

Having delineated the common core, we next sought to generate a phylogenetic tree relating diverse GT-A fold families using the core alignment. Because of the inherent challenges in the generation and visualization of large trees (19), we used a representative set of GT-A fold sequences for phylogenetic analysis by first clustering the ~600,000 sequences into functional categories using a Bayesian Partitioning with Pattern Selection (BPPS) method (20). The BPPS method partitions sequences in a multiple sequence alignment into hierarchical sub-groups based on correlated residue patterns characteristic of each sub-group (Methods). This revealed 99 sub-groups with distinctive patterns. Representative sequences across diverse phyla from these sub-groups (993 sequences, Dataset S2) were then used to generate a phylogenetic tree (Fig. 2). Based on the phylogenetic placement of these sequences, we broadly define fifty-three major sub-groups, thirty-one of which correspond to CAZy-defined families (Table S2). The remaining sub-groups correspond to sub-families within larger CAZy families. In particular, we sub-classified the largest GT family in the CAZy database, GT2, into ten phylogenetically distinct sub-families. Likewise, GT8 and GT31 were classified into seven and five sub-families, respectively. These sub-families are not explicitly captured in CAZy and are annotated based on overall sequence similarity to functionally characterized members. For example, “GT2-LpsRelated” corresponds to a sub-family within GT2 most closely related to the bacterial β-1-4-glucosyltransferases (lgtF) involved in Lipopolysaccharide biosynthesis (Fig. 2, Fig. S4). Such a hierarchical classification captures the evolutionary relationships between GT-A fold families/sub-families while keeping the nomenclature consistent with CAZy.

**Figure 2:**
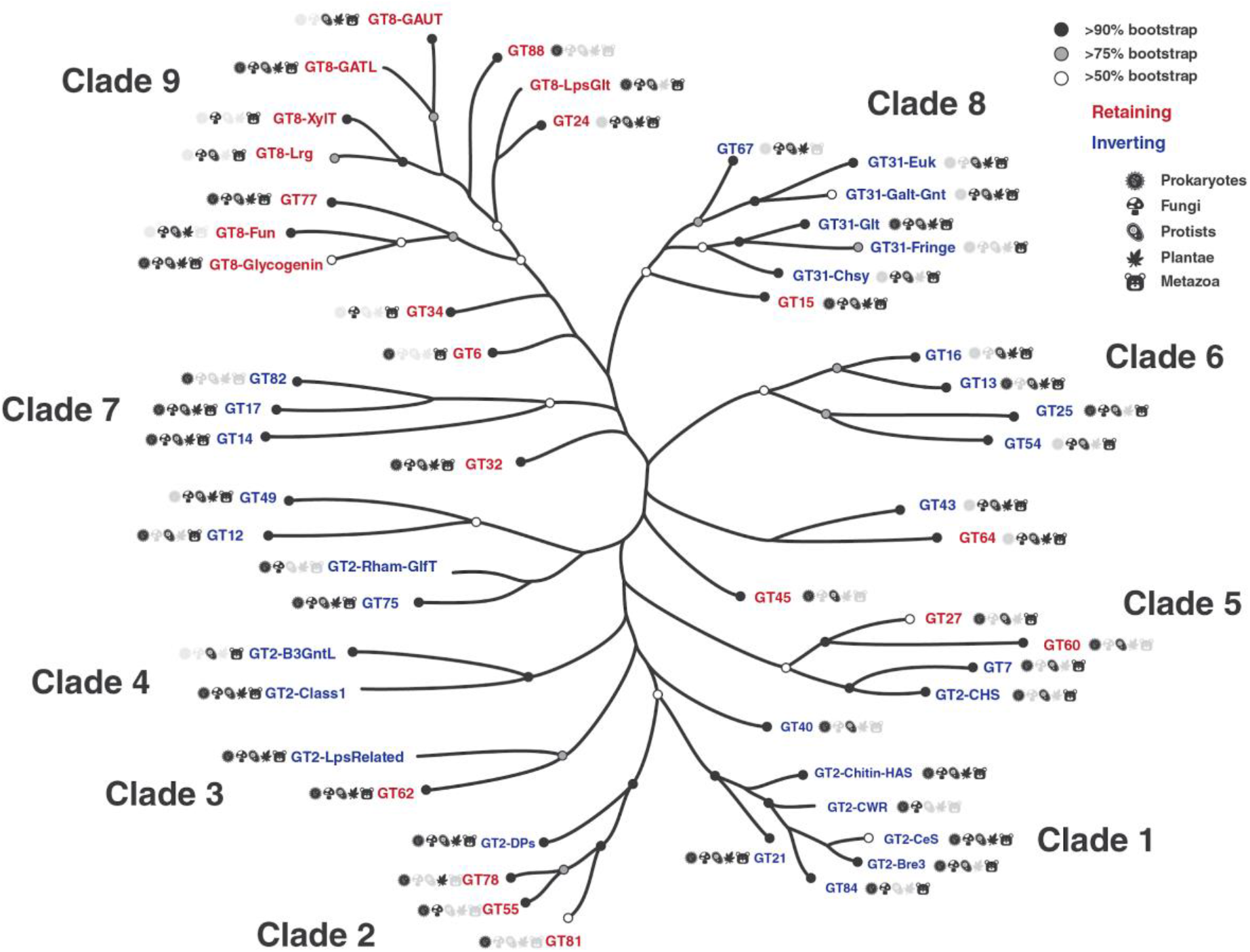
Phylogenetic tree highlighting the 53 major GT-A fold subfamilies. Tips in this tree represent GT-A sub-families condensed from the original tree for illustration. Support values are indicated using different circles. Circles at the tips indicate bootstrap support for the GT-A family clade represented by that tip. Tips missing the circles represent GT-A families that do not form a single monophyletic clade. Nodes missing circles have a bootstrap support less than 50% and are unresolved. Icon labels indicate the taxonomic diversity of that sub clade. Colors indicate the mechanism for the families (blue: Inverting, red: Retaining). Detailed tree with support values and expanded nodes are provided in Fig. S4 and in Newick format in Dataset S4. The family names are described in Table S2.

GT-A fold families and sub-families can be further grouped into clades based on shared sequence features and placement in the phylogenetic tree (Fig. 2). For example, clade 1 places four GT2 sub-families (GT2-CeS, GT2-CWR, GT2-Chitin-HAS and GT2-Bre3) with GT84 and GT21 supported by high bootstrap values. Members of these six families are all involved in either polysaccharide or glycosphingolipid biosynthesis. Additionally, the pattern-based classification identified a conserved [QR]XXRW motif in the C-terminal HV3 (Fig. S5) which is unique to members of this clade. The [QR]XXRW motif residues coordinate with the donor and acceptor in a bacterial cellulose synthase (from GT2-CeS family) (21) and mutation of these residues in bacterial cyclic β-1,2-glucan synthetase (Cgs, GT84) abrogates activity (22), suggesting a critical role of this motif in functional specialization of clade 1 GT-As.

The GT8 sub-families form sub-clades within the larger clade 9. For example, GT8 sequences involved in the biosynthesis of pectin components group together in the GT8-GAUT and GT8-GATL families (Fig. 2). The human LARGE1 and LARGE2 glycosyltransferases are multi-domain enzymes with two tandem GT-A domains. Their N-terminal GT-A domains fall into the GT8-Lrg subfamily that groups closely with GT8-xylosyltransferase (GT8-XylT) subfamily enzymes and places all the GT8 xylosyltransferases into a single well supported sub clade. The lipopolysaccharide α-glucosyltransferases (GT8-LpsGlt) group with the glucosyltransferases of the GT24 family, suggesting a common ancestor associated with glucose donor specificity. On the other hand, the GT8-Glycogenin sub family, which also includes members that transfer a glucose, is placed in a separate sub-clade, possibly indicating an early divergence for its unique ability to add glucose units to itself (23). Clade 9 members also share common sequence features associated with substrate binding that includes a lysine residue within the commonly shared KPW motif in HV3 that coordinates with the phosphate group of the donor (e.g. bacterial LgtC (GT8-LpsGlt) and other structures of clade 9 members)(Fig. S5).

We noticed that three out of four MGAT GT-A families responsible for the branching of N-glycans (GT13 MGAT1, GT16 MGAT2 and GT54 MGAT4) fall in the same clade (clade 6), as expected (Fig. 2). In contrast, the fourth family, GT17 MGAT3, which adds a bisecting GlcNAc to a core β-mannose with a β-1,4 linkage, is placed in a separate clade with GT14 and GT82 (clade 7), while a fifth MGAT member creating β-1,6-GlcNAc linkages (GT18 MGAT5) is a GT-B fold enzyme (24).

We further note that fifteen out of fifty-three GT-A families are found in both prokaryotes and eukaryotes. These fifteen families fall on different clades throughout the tree. GT-A families present only in prokaryotes, like GT81, GT82 and GT88, are also spread out in different clades (Fig. 2). Similarly, other GT-A families that are present within restricted subsets of taxonomic groups (like GT40 and GT60 present only in prokaryotes and protists) are also scattered throughout the tree. These observations suggest that the divergence of most GT-A families predates the separation of prokaryotes and eukaryotes.

### Multiple evolutionary lineages for inverting and retaining mechanisms

To obtain insights into the evolution of catalytic mechanism, we annotated the phylogenetic tree based on known mechanisms of action (inverting or retaining). Inverting GTs are colored in blue in the phylogenetic tree, while retaining GTs are colored in red (Fig. 2). The dispersion of inverting and retaining families in multiple clades suggests that these catalytic mechanisms emerged independently multiple times during GT-A fold evolution. We find that natural perturbations in the catalytic base residue, an important distinction between the inverting and retaining mechanisms, correlates well with these multiple emergences across the tree. The residue that acts as a catalytic base for inverting GTs (aspartate within the xED motif, xED-Asp) is variable across the retaining families consistent with its lack of role in the retaining SNi mechanism (6). In the inverting families, the xED-Asp is nearly always conserved and appropriately positioned to function as a catalytic base (Fig. 3A), though some exceptions have been noted (6, 25). Out of the five clades grouping inverting and retaining families, inverting families in three of these clades do not conserve the xED-Asp (GT2-DPs, GT2-LpsRelated and GT43). The heterogeneous nature of this residue in these families suggests that change of the catalytic base residue could be a key event in the transition between inverting and retaining mechanisms. Unlike families that conserve the xED-Asp, these families achieve inversion of stereochemistry through alternative modes that may relieve the constraints necessary to conserve the xED-Asp. For example, in GT43, the Asp base is replaced by a glutamate residue, which shifts the reaction center by one carbon bond (6). Further, the dolichol phosphate transferases (DPMs and DPGs) in the GT2-DP family, which lack the xED-Asp entirely, transfer sugars to a negatively charged acceptor substrate (a phosphate group) and thus do not need a catalytic base to initiate nucleophilic attack (25). Other GT-A inverting families lacking the xED-Asp (GT12, GT14, GT17, GT49 and GT82) are grouped into separate monophyletic clades segregating them from inverting families with conserved the xED-Asp (Fig. 2). Out of these, only GT14 has representative crystal structures where a glutamate serves as the catalytic base (26). For other inverting families with a non-conserved xED-Asp, residues from other structural regions may serve as a catalytic base. On the other hand, retaining families like GT64 conserve the xED-Asp, yet do not use it as a catalytic base. Thus, there may be multiple ways in which inverting and retaining mechanisms diverge, with one path being mutation of this xED-Asp catalytic base.

**Figure 3:**
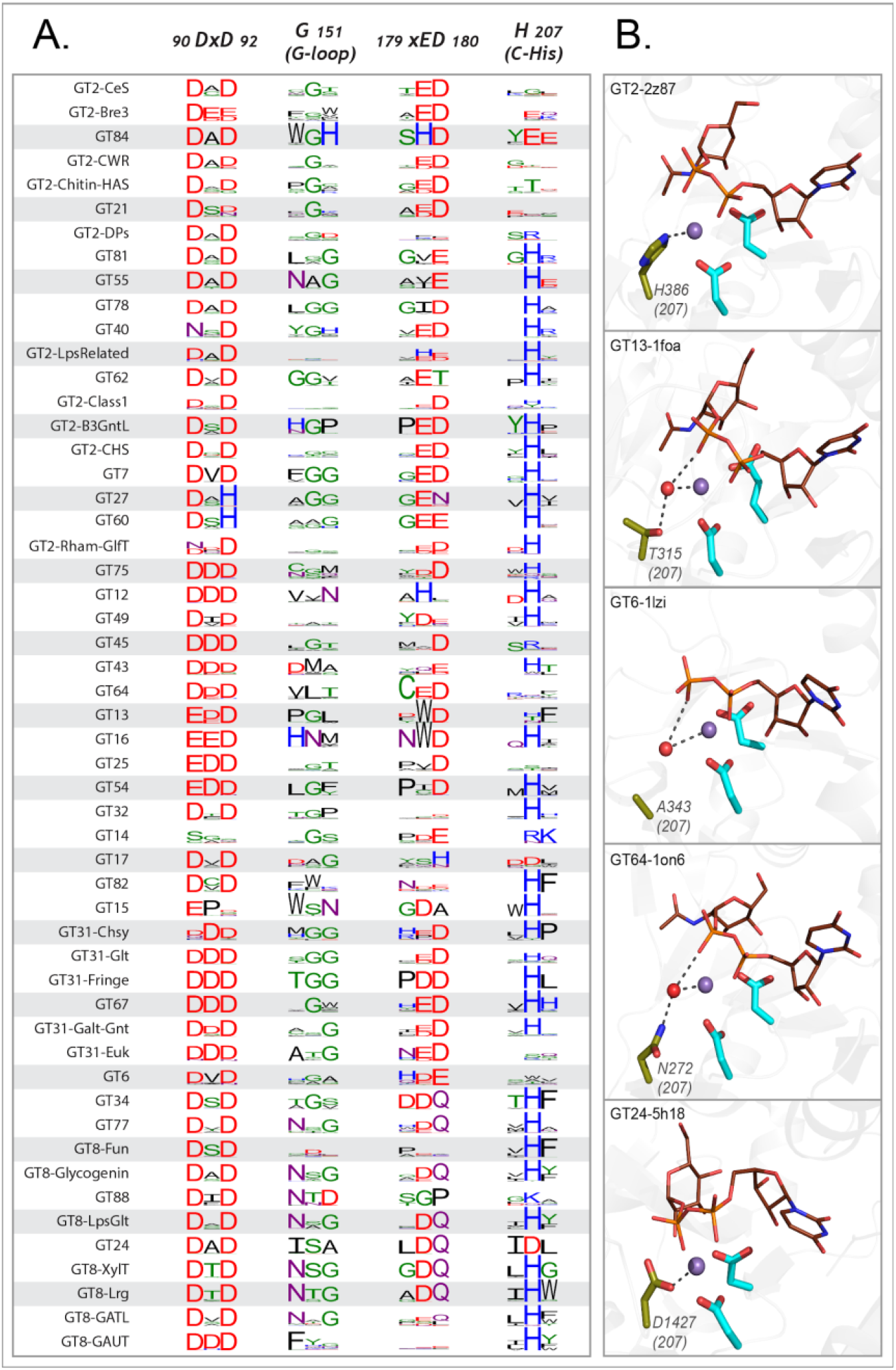
Variations in the conserved core of the GT-A families. A) Weblogo depicting the conservation of active site residues in the common core are shown for each of the GT-A families. Residues are colored based on their physiochemical properties. B) Variations in the C-His is compensated either using a water molecule (red sphere) or other charged residues (olive sticks) to conserve its interactions. The metal ion is shown as a purple sphere. The donor substrate is shown as brown lines. Interactions between the residues, metal ion and the donor are shown using dotted lines.

One strongly supported clade that includes both inverting and retaining families is clade 2 that groups inverting GT-A family members that transfer sugars to phosphate acceptors (GT2-DPs) with three retaining GT-A families that also have phosphate-linked acceptors (GT55, GT78 and GT81). This placement is further supported by the observation that these families share structurally equivalent conserved residues in the HV2 region that coordinate the phosphate group of the acceptor. In the GT2-DP subfamily, R117, R131 and S135 (Fig. 4A) in HV2 coordinate with the acceptor phosphate groups. The conservation of these residues in GT55 and GT81 suggests that they likely perform similar interactions in these latter subclades. Indeed, in the crystal structure of M. tuberculosis GpgS (GT81), HV2 adopts a conformation similar to GT2-DPs and the shared residues G184, R185 and T187 (equivalent to R117, R131 and S135) form similar interactions with the phosphate group of the acceptor (Fig. S5).

**Figure 4:**
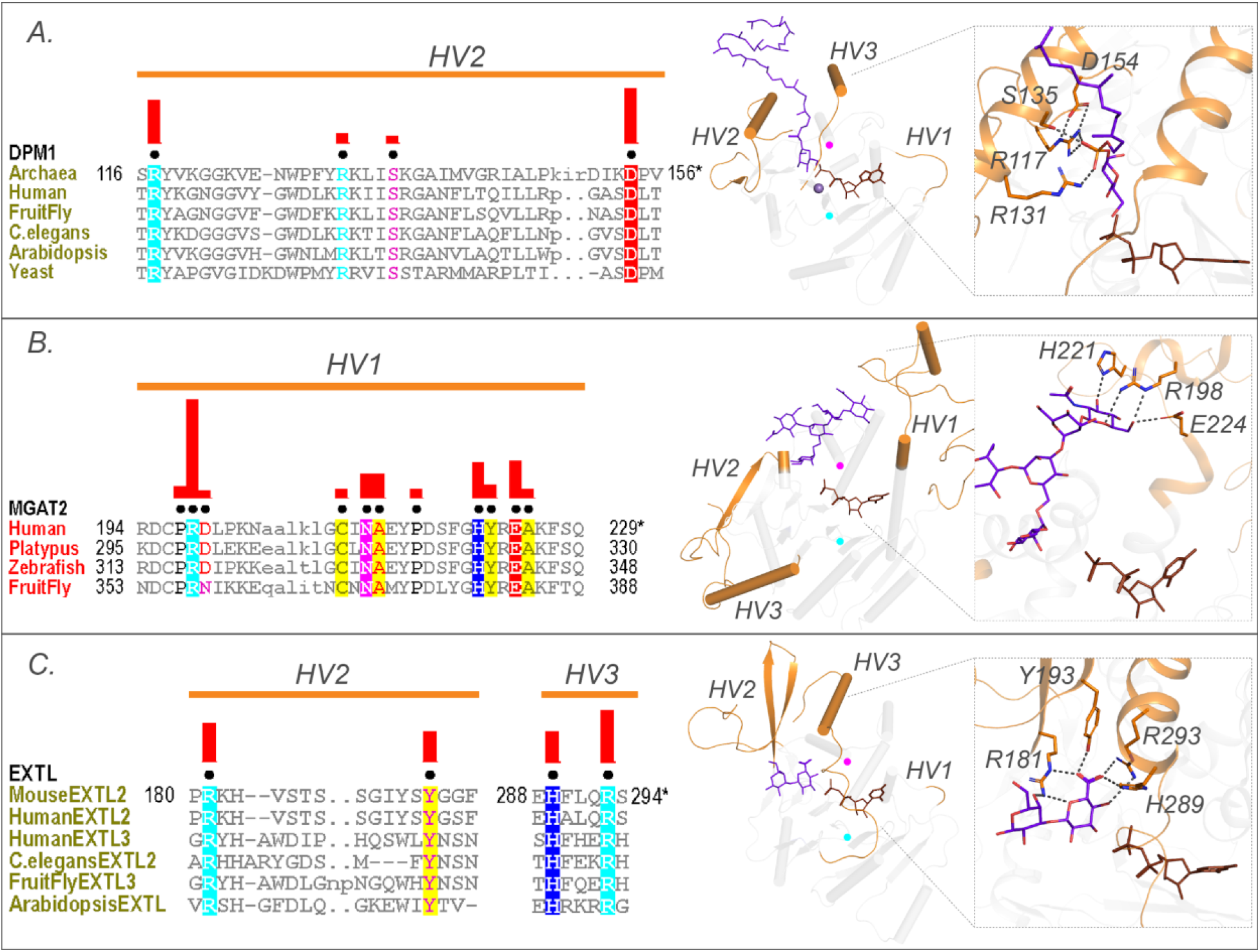
Family specific conserved features in the HV regions help coordinate the acceptor. Conserved residues in A) HV2 of the DPM1 sequences in the GT2-DP subfamily coordinate the phosphate group of the acceptor. B) HV1 of GT16 MGAT1 provide acceptor specificity. C) HV2 and HV3 of EXTL GT64 family (C-terminal GT domain of the multidomain sequences) coordinate the acceptor. Left: Alignments highlighting the constrained residues are shown for each family. The family specific conserved residues are shown using black dots above the alignment. Red bars above these dots indicate the significance of conservation (Higher bar corresponds to more significantly conserved position). Right: Representative pdb structures are shown for each family (GT2-DP:5mm1, GT16:5vcs, GT64:1on8); Donor substrates are colored brown. Acceptors are colored purple. HVs are highlighted in orange. The position of the conserved DxD and xED motif for each structure is shown as cyan and magenta circles respectively.

Clade 5 places the inverting GT7 and GT2-CHS with the retaining GT27 and GT60 families (Fig. 2). This supports the evolution of these families from a close common ancestor through gene duplication and divergence, which has been suggested through structural similarities between GT7 and GT27 (27). After this initial divergence in mechanism within clade 5, the subclades group the β-1,4-GalNAc transferase domains of bacterial and protist chondroitin polymerases (involved in the elongation of glycosaminoglycan chondroitin)(GT2-CHS) with the GT7 family. The GT7 family includes the higher organism counterparts of the β-1,4-GalNAc transferase domains of chondroitin synthases, along with β-1,4 Gal transferases. The close placement of GT60 and GT27 families in this clade is also directly supported by previous literature indicating that these families share a conserved mode of polypeptide Ser/Thr O-glycosylation (28). Clade 5 thus consolidates previous independent findings and suggests a shared ancestor, potentially extending the common ancestry of GT2-CHS and GT7 to include GT27 and GT60, with an ancestral divergence in mechanism.

### Variations in the core and hypervariable regions contribute to unique modes of substrate specificity

Analysis of the patterns of conservation and variation in the common core indicates that each residue position within the core has been mutated in some context during the course of evolution, highlighting the tolerance of the GT-A fold to extensive sequence variation. While some of these variations are confined to specific clades or families, such as replacement of DxD motif with DxH motif in GT27 and GT60, other variations are found independently across distal clades (Fig. 3A). For example, GT14 and prokaryotic members of GT6 that fall on different clades, have independently lost the DxD motif and no longer require a metal ion for activity (26, 29).

The C-His is also lost independently in multiple clades (Fig.3A). In order to investigate how the loss of metal binding C-His is compensated, we analyzed the C-His-metal ion interactions across all available crystal structures. Structural alignment of crystal structures from families that are missing the C-His such as GT13, GT6 and GT64 families revealed a water molecule coordinating the metal ion in a manner similar to the C-His sidechain (Fig. 3B). In other families, such as GT24, we found that the C-His is substituted by an aspartate (D1427), which coordinates with the metal ion similar to C-His (Fig. 3B, bottom panel). Likewise, the conserved hydrophobic coupling between αF helix and the Rossmann domain is replaced by charged interactions (R388 and E274, respectively) in some retaining GTs such as GT15 and GT55 (Fig. S2). These substitutions point to the ability of GT-As to accommodate changes, even in conserved positions at the core, through compensatory mechanisms.

As noted above, we found that the hypervariable regions display significant variations across GT families but conserve family specific residues that contribute to acceptor specificity. For example, a distinctive arginine (R117) and aspartate (D154) along with R131 and serine S135 within the HV2 of DPM1 (GT2-DP sub-family) contribute to specificity towards a dolichol phosphate acceptor by creating a charged binding pocket for the phosphate group (Fig. 4A). Likewise, family-specific residues (R198, H221 and E224 in 5vcm) within the HV1 of MGAT2 (GT16) form a unique scaffold for recognizing the terminal GlcNAc of the N-glycan acceptor (Fig. 4B). Similarly, the C-terminal GT64 domain of the multidomain EXTLs contain specific residues in HV2 (R181 and Y193) and HV3 (H289 and R293) that form a unique binding pocket for the tetrasaccharide linker acceptor used to synthesize glycosaminoglycans (Fig. 4C). Together these examples illustrate the ability of HVs to evolve family specific motifs to recognize different acceptors.

### Machine learning to predict the donor specificity of GT-A sequences

As discussed above, the conserved catalytic residues dictate the mechanism of sugar transfer and metal binding while the extended HVs use family specific motifs to dictate acceptor specificity. We also find some clade specific features (such as the conserved Lys in clade 9, and QXXRW in clade 1) and G-loop residues involved in donor binding, however, the overall framework that dictates donor sugar specificity in GTs is largely unknown. Sequence homology alone is insufficient to predict donor specificity because evolutionarily divergent families can bind to common substrates, and sometimes even two closely related sequences bind to different donors (Fig. S6)(15). Our global analysis of GT-A families provides a comparative basis to contrast sequences that bind to different donors. To test whether evolutionary features gleaned from this global analysis can be used to better predict donor substrate specificity, we employed a machine learning (ML) framework that learns from the specificity-determining residues of functionally characterized enzymes to predict specificity of understudied sequences. In brief, using an alignment of a well curated set of 713 GT-A sequences (Dataset S5, SI Methods) with known donor sugars, we derived five amino acid properties (hydrophobicity, polarity, charge, side chain volume and accessible surface area) from each aligned position within the common core. These properties were then used as features to train multiple machine learning methods. Among the six methods used, random forest model achieved the best prediction performance (accuracy ~88%) based on a 10-fold cross validation (CV) using 239 contributing features (SI Methods, Fig. 5A,B, Table S6). To further validate the model, we tested its performance on a validation set of 64 sequences that were not used to train the ML model but have known sugar specificities. The random forest classifier correctly predicted donor substrates for 92% of these sequences, nearly 80% of which were predicted with high confidence (blue rows in Dataset S6).

**Figure 5:**
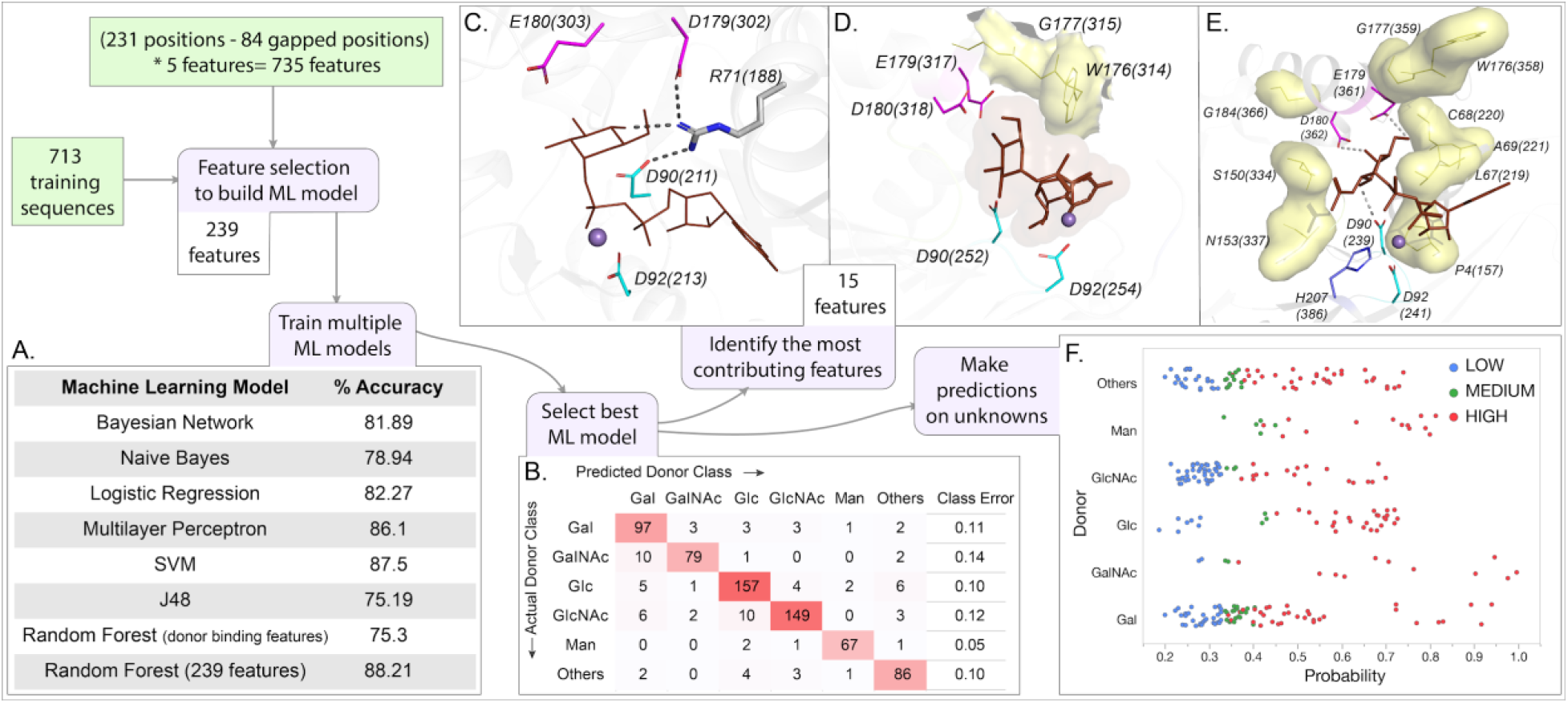
Outline and results of the ML analysis. Training set input into the pipeline are shown in green squares. Steps of the ML analysis in purple boxes are associated with different panels of the figure. A) Percent accuracy based on 10-fold CV for each of the trained ML models. B) Confusion matrix from the best model (random forest using 239 features). The full model was used to generate this matrix which was then evaluated using 10-fold CV. Panels C, D and E show the 15 most contributing features mapped into representative structures. Metal ion and donor substrate are shown in purple sphere and brown sticks respectively. C) R71 interacts with the donor sugar and forms a bridge between D179 and D90 in GT6 (5c4b). D) The top contributing features (position 176 and 177, yellow surface) line the donor binding pocket in GT7 (1o23). E) The top contributing feature positions not directly involved in donor binding fall around the active site and are shown using yellow surface in the prototypic GT-A (2z87). F) Scatter plot showing the probability scores assigned for each predicted sequence by the predicted donor type. Colors indicate the confidence level of the prediction derived using the probability and its difference from the 2nd class (SI Methods, Dataset S6)

This model was then used to predict donor sugars for GT-A domains with unknown specificities from 5 organisms: *H. sapiens*, *C. elegans*, *D. melanogaster*, *A. thaliana* and *S.cerevisiae* (Dataset S6). Each prediction is associated with a confidence level derived from the probability for each of the 6 donor classes (SI Methods). 55% of the predictions have high and moderate confidence levels and present good candidates for further investigation (Fig. 5F). The remaining 45% of the predictions are low confidence. This likely reflects the promiscuity of GT-As for donor preferences, as seen across many GT-As (16, 30), or non-catalytic GT-As like C1GALT1C1 (Cosmc) (31).

Our predictions assign putative donors for 10 uncharacterized human GT-A domains (Dataset S6). B3GNT9 is predicted to employ UDP-GlcNAc with high confidence like other GT31 β-3-N-acetylglucosaminyltransferases (B3GNTs) in humans (32). The two procollagen galactosyltransferases in humans (COLGALT1 and COLGALT2) are multidomain proteins with two tandem GT domains. While their respective C-terminal domains catalyze β-Gal addition to hydroxylysine side-chains in collagen, our predictions assign a putative GlcNAc transferase role for their N-terminal GT domain. More interestingly, GLT8D1, a GT8 glycosyltransferase with an unknown function implicated in neurodegenerative diseases (33), is predicted to have a glucosyltransferase specificity. In other organisms, the GT2 sequences in *A. thaliana* (mostly involved in plant cell wall biosynthesis) are predicted to bind glucose and mannose substrates, the primary components of the plant cell wall (Dataset S6). We also identify a novel galactosyltransferase function for a GT25 enzyme in *C. elegans*. These predictions can guide characterization of new GT sequences with unknown functions.

We next performed feature selection to identify features that contribute most to substrate (donor) prediction. Fifteen features selected by a combination of multiple feature selection methods (SI Methods) contributed most to substrate prediction. Some of these features correspond to residues involved in substrate binding and catalytic functions such as the Asp within the DxD motif, residues in the G-loop and the C-His (15, 16, 25). One such contributing feature is a positively charged residue at aligned position 71 that emanates from the αC helix and interacts with the donor sugar. In a crystal structure of ABO (GT6), R71 (R188 in 5c4b) has been shown to bridge the DxD and the xED motif to keep the catalytic site intact (Fig. 5C)(34). Our ML model identifies the charge and accessible surface area of R71 as a major contributing feature in donor specificity. The remaining features, surprisingly, are not directly involved in donor binding. For example, the total volume of the residues in a loop preceding the xED motif (aligned position 176-177, WG358-359 in 2z87, Fig. 5D) contributes significantly to donor specificity (Table S3), presumably by controlling the accessibility of the donor binding site. Further, an ML model trained using features from only the donor binding residues performs with an accuracy of only 75%, indicating the importance of features other than those directly involved in donor binding. Thus, despite only a few residues being directly involved in donor interactions, additional contributions to donor specificity come from residues more distal from the active site. Contributions from these peripheral secondary shell features surrounding the donor binding site (Fig. 5E) highlight the potential role of higher order (allosteric) interactions in determining donor substrate specificity.

## Discussion

Prior studies on the evolution of GTs have generally focused either on distinct GT subfamilies or biosynthetic pathways with additional structural classifications of GTs into one of three distinct protein fold superfamilies (6, 9, 10). In our present work we focused on the analysis of the largest of the GT superfamilies, those that comprise a GT-A protein fold characterized by an extended Rossmann domain with associated conserved helical segments. These enzymes generally employ the Rossmann domain for nucleotide sugar donor interactions and extended loop regions for acceptor glycan interactions (6). Using an unbiased profile search strategy, we assembled a total of over 600,000 GT-A fold related sequences from all domains of life for deep evolutionary analysis. To support this profile-based assembly, we leveraged structural alignments on GT-A fold enzymes in PDB and secondary structure predictions when no crystal structures were available. The resulting alignment allowed the definition of a common structural core shared among the diverse GT-A fold enzymes and defined positions where hypervariable loop insertions were elaborated to provide additional functional diversification (Fig. 1). In cases where data was available for enzyme-acceptor complexes these latter loop insertions generally contribute to unique, family specific acceptor interactions. Thus, a structural framework is presented for GT-A fold enzyme evolution. Since the common core is present across all kingdoms of life, it presumably represents the minimal ancestral structural unit for GT-A fold catalytic function by defining donor substrate interactions and minimal elements for acceptor recognition and catalysis. In fact, we find several archaeal and bacterial sequences that closely resemble this common core consensus sequence (Dataset S7). Based on our studies, we propose a progressive diversification of glycosyltransferase function through evolution of donor specificity by accumulation of mutations in the common core region and divergence in acceptor recognition through expansion of the hypervariable loop regions. Consistent with this view, we find conserved family-specific motifs within the hypervariable regions that confer unique acceptor specificities in various families. These expansions likely contributed to the evolution of new GT functions and catalyzed new glycan diversification observed in all domains of life.

A surprising finding from our studies is the dispersion of inverting and retaining catalytic mechanisms among families in the GT-A fold evolutionary tree (Fig. 2). Recent models indicate that distinctions between inverting and retaining catalytic mechanisms arise from differences in the angle of nucleophilic attack by the acceptor toward the anomeric center of the donor sugar (6). Inverting mechanisms require an in-line attack and direct displacement by the nucleophile relative to the departing nucleotide diphosphate of the sugar donor and a conserved placement of the xED-Asp carboxyl group as catalytic base at the beginning of the αF helix. In contrast, retaining enzymes generally alter the angle of nucleophilic attack by the acceptor, use a donor phosphate oxygen as catalytic base, and employ a dissociative mechanism for sugar transfer (6). The fundamental differences in these catalytic strategies would suggest an early divergence of enzymes employing these respective mechanisms. However, the GT-A fold phylogenetic tree strongly suggests that inverting and retaining mechanisms evolved independently at multiple points in the evolution of GT-A families (Fig. 2). Since the main difference in these mechanisms is the change in position of the nucleophilic hydroxyl and catalytic base, we believe this poses the possibility for a transitional phase in the evolution between the two mechanisms. The xED-Asp carboxyl group is highly conserved in the inverting enzymes and is appropriately placed for acceptor deprotonation. Variants of this motif either lack the residue entirely, as seen in many retaining enzymes, or use compensatory modes to accommodate changes at this position, as seen for the inverting enzymes in GT43, GT2-DPs, and GT2-LPSRelated. In fact, in each of the latter cases the respective inverting GT family is clustered with closely related GT families employing a retaining catalytic mechanism. Thus, inverting enzyme variants that accommodate changes to the xED motif group may represent examples of transitional phases in evolution between inverting and retaining catalytic mechanisms. Other inverting enzymes harboring variants in the xED motif segregate into separate clades and could represent outlier families that have developed alternative ways to compensate for the loss of xED-Asp. This ability to evolve distinct catalytic strategies, in some cases through presumed convergent evolution, could allow each family to evolve independent capabilities for donor and acceptor interactions as well as for anomeric linkage of sugar transfer, while retaining other essential aspects of protein structural integrity through the use of a conserved and stable Rossmann fold core.

In an effort to define the sequence constraints for the respective catalytic mechanisms we also employed a machine learning framework for prediction of the mechanism for unknown sequences and were able to assign the donor sugar nucleotide for a test set of enzymes with high accuracy. Surprisingly, the contributing features for accurate prediction include residues involved in donor binding as well as positions that are distal to the active site likely as secondary shell effects or allosteric interactions. Due to their indirect involvement, such positions are generally difficult to pinpoint using structural studies alone emphasizing the need for robust sequence-based comparative analysis to understand GT-A function. The predictions made from the ML framework can serve as a valuable resource for generating and testing new hypotheses on GT-A functions.

Numerous additional insights into GT function were also revealed through inspection of the aligned sequences and the phylogenetic tree. For example, the clustering of mammalian N-glycan GlcNAc branching enzymes (MGAT1 (GT13), MGAT2 (GT16), and MGAT4 (GT54)) in the same clade suggests a common origin for these enzymes, while placement of MGAT3 (GT17) in a separate clade could point to its unique role in adding a bisecting GlcNAc to the N-glycan core thereby regulating N-glycan extension (35). In contrast, MGAT5 (GT18) involved in N-glycan β1,6-GlcNAc branching is a GT-B fold enzyme with a clearly distinct evolutionary origin. While most clades are well resolved, bootstrap support values for nodes at the base of the tree are low and need to be interpreted with caution. This low resolution results from high divergence between families and possibly other events like horizontal gene transfer and convergent evolution. However, trees generated using alternative strategies support the overall topology (Fig S7) and clades are congruent with clusters obtained using an orthogonal Bayesian classification scheme, which adds confidence to the phylogeny (Table S2).

For some GT-A fold enzymes variations in the catalytic site can also be accommodated by other compensatory changes. An example is the use of the C-His motif for coordination of the divalent cation in most GT-A fold enzymes in contrast with enzyme variants that employ water molecules to compensate for the loss of this residue (Fig. 3B). Similarly, some inverting GTs dispense with the use of the divalent cation and the DxD motif and substitute interactions with the sugar donor through use of basic side chains (e.g. GT14). A further extreme is the duplication, divergence and pseudogenization within the GT31 family. Human C1GALT1C1 (GT31, COSMC) shares a high sequence similarity to another GT31 member, C1GALT1 (T-synthase), yet COSMC has lost both the DxD and the xED motifs and has no catalytic activity. Instead, COSMC acts as an important scaffold and chaperone for the proper assembly and catalytic function of T-synthase (31). The ability of GT-As to harbor such structural variations that allow them to develop new functions make them well-suited to evolve rapidly and facilitate the synthesis of a diverse repertoire of glycans across all living organisms.

Our unbiased, top-down sequence-based analysis suggests new and unanticipated evolutionary relationships among the GT-A fold enzymes. Prior suggestions of such relationships have been inferred by the clustering of GT sequences into families in the CAZy database. However, the CAZy database of GT sequences does not provide access to the broader sequence relationships among the GT-A fold enzymes or how a general model of a core conserved GT-A fold scaffold can serve as a progenitor catalytic platform for binding sugar donors and facilitating glycan extension. The sequence assembly, phylogenetic tree, and placement within the framework of known GT-A fold structures in the present studies provide key insights into conserved elements of the hydrophobic core, linkage to the DxD motif for cation and sugar donor interactions, and the conserved αF helix harboring the xED catalytic base. Additional hypervariable extensions at defined positions from this conserved core were then progressively recruited to confer unique modes of acceptor interactions to develop new specificities and evolve new functions. Thus, the core of the protein scaffold can be maintained to facilitate protein stability while rapid evolution of the hypervariable loops can develop new glycan synthetic functionalities through presentation of novel acceptors to the catalytic site. Variation in the location of the acceptor hydroxyl nucleophile relative to the donor sugar anomeric center presents the opportunity for distinctions in catalytic mechanism and anomeric outcome for sugar transfer. The result is a rapidly evolving set of GT enzymatic templates as the biosynthetic machinery for diverse glycan extension on cell surface and secreted glycoproteins and glycolipids. In such contexts the resulting glycoconjugates confer potential functional selective advantages at the cell surface, but also act as ligands and pathogen entry points for negative evolutionary pressure. The constant challenges to adapt to these Red Queen effects of positive and negative selective pressures for glycan synthesis have led to the remarkable diversity in the GT enzymes and their resulting glycan structural products. We anticipate that the sequence and structural principles that drive GT-A fold evolution will also likely extend to GT-B and GT-C fold enzymes and represent a common theme for the elaboration of diverse glycan structures in all domains of life.

## Methods

### Generation of GT-A profiles and alignment

Multiple alignments for 34 CAZy GT-A were collected from the Conserved Domain Database (CDD) (36) or were manually built using MAFFT v7.3 (37) from sequences curated at the CAZy database (Table S1). These seed profiles were then multiply aligned using the mapgaps scheme (14) guided by a structure based sequence alignments of all available pdb structures using Expresso (38) and MAFFT to generate the GT-A profiles. Representative pdb structures described in this study are listed and cited in Dataset S1. Finally, the alignment of secondary structures and conserved motifs were manually examined and corrected, where necessary. Very divergent GT-A families such as GT29 and GT42 sialyltransfearses were not included in this analysis (SI Methods). The GT-A profiles were then used for a sequence similarity search using mapgaps to identify and align more than 600,000 GT-A domain sequences from the NCBInr database. This alignment was filtered for fragmentary sequences and false hits. This filtered alignment was then used to define the boundaries of the GT-A common core (SI Methods).

### Bayesian Statistical analyses

A representative subset of 24,650 GT-A sequences were generated from the ~600,000 putative GT-A sequences by using a family-based sequence similarity filtering (SI Methods). This sequence set was then used to apply the Optimal multiple-category Bayesian Partitioning with Pattern Selection (omcBPPS) scheme (20). omcBPPS identifies patterns of column-wise amino acid conservation and variation in the multiple sequence alignment. The resulting family specific positions were then used as statistical measures to classify the GT-As into 99 unique sets that correspond to the 53 families described in this study (Table S2). omcBPPS also identified aligned positions that are conserved across all GT-A fold families. This revealed the 20 conserved positions within the core component, that were also verified by calculating conservation scores using the Jensen-Shannon divergence score as described and implemented by (39)(used in Fig. 1A).

### Phylogenetic analysis

A smaller subset of 993 sequences were used for phylogenetic reconstruction of the GT-A families. This set includes all the identified GT-A sequences from five model organisms: *H. sapiens* (human), *C. elegans* (worm), *D. melanogaster* (fly), *A. thaliana* (dicot plant) and *S. cerevisiae* (yeast) along with select sequences representing the diverse taxonomic group in each family. These representative sequences were selected by finding the union of top hits for every taxonomic group present within each of the 99 sets and the seed alignments for the 34 CAZy GT-A families. This selection criteria maximized the phylogenetic and taxonomic diversity while keeping the number of sequences to a minimum. The alignment for these 993 sequences were then trimmed to remove the insert positions and keep only the 231 aligned positions described above. This trimmed alignment was used to build a phylogenetic consensus tree using IQTree v1.6.1 (40) with the following options: -nt AUTO -st AA -m MFP+MERGE -alrt 1000 -bb 1000 -wbt -nm 1000 -bnni. Further support for the phylogenetic tree was collected by comparing its topology to trees generated using orthogonal methods like Hidden Markov Model (HMM) distances and structural similarities, that have been used in previous studies (41, 42)(Fig. S7, SI Methods).

### Defining the GT-A families and sub-families

The GT-A sequences were first classified into pattern-based groups using omcBPPS. Based on the placement of representative sequences from these groups in the phylogenetic tree, they were merged into GT-A families and sub-families. The correspondence between the 53 GT-A families and subfamilies with the 99 pattern-based groups are provided in Table S2. Sequences from some families did not form any distinct pattern-based groups due to either a low number of sequences for a statistically significant grouping (GT78) or a lack of distinguishing patterns within the aligned positions (GT25, GT88). Representative sequences for these families were collected from the seed alignments for these families as described above. We also identified the N-terminal GT2 domain of the multidomain chondroitin polymerase structure from E. coli (Pdb Id: 2z87) as the prototypic GT-A structure to use as a comparative basis for structural analyses. This sequence was selected based on the lowest E-value and highest similarity score of a BLAST search of all pdb structures against the GT-A consensus sequences. Weblogos for the conserved active site residues were derived for each GT-A subfamily using Weblogo 3.6.0 (43).

### Machine learning analysis

In order to train an ML model for GT-A donor substrate prediction, we first curated a training dataset by mining the “characterized” tab of the CAZy GT database and the UniProt database to find 713 GT-A domain sequences with known donor sugars. The donor sugar information for these sequences were extracted from their assigned protein names. Based on the availability of training sequences, 6 major donor type classes were defined: Glc, GlcNAc, Gal, GalNAc, Man, and “Others” with each class having more than 70 sequences in the training dataset. The “Others” category merged the least represented donor types with less than 50 training sequences each (Ara, Fuc, GalF, GlcA, ManNAc, Rham, and Xyl). An alignment of the 713 sequences were generated which was filtered and then used to derive 5 amino acid properties (charge, polarity, hydrophobicity, average accessible surface area, and side chain volume) for each aligned position that were used as features for machine learning. We implemented correlation-based feature selection (CFS) (44) with 5-fold CV by using WEKA version 3.8.3 (45) under default settings to select 239 informative features for building multiple multiclass classification models.

Using these features, we trained multiple models (SVM, multilayer perceptron, Bayesian network, logistic regression, naive Bayes classifier, J48, and random forest) using WEKA and the R package “randomForest” (46). These models were compared using 10-fold CV under default settings. 10-fold CV evaluates the ML models by iteratively training on 90% of the data selected at random and testing the prediction on the unseen 10% of the data. This is repeated 10 times and the results on the testing dataset are summarized into an accuracy measure. The random forest model trained with 239 features had the highest accuracy and overall performance and thus was selected as the model of choice for predicting donor sugar substrates for GT-A enzymes. Confidence scores were assigned for each prediction based on the probability for each of the 6 donor classes. Further details of the methods implemented for machine learning and generation of confidence levels are provided in SI Methods.

## Author Contributions

R.T., A.S.E., K.W.M. and N.K. designed research; R.T., A.V., L.C.H. and W.Y. performed research; R.T., A.V., L.C.H., K.R., K.W.M. and N.K. analyzed data; and R.T., A.V., K.W.M. and N.K. wrote the paper.

