## Supplementary Information for "Deep evolutionary analysis reveals the design principles of fold A glycosyltransferases"

### SI Methods

#### Building the GT-A profile alignments

We created a shortlist of all known and putative GT-A fold families in the CAZy database to be included in this analysis. To do this, we first included the 17 families annotated as having a GT-A fold in the CAZy database. Then, based on published literature (1–3), we added 20 more families that were putatively listed as being the GT-A fold. For each of these families, representative alignments were collected from the Conserved Domain Database (CDD) where available. For families where a CDD alignment was not available, we selected representative sequences from the CAZy database and aligned them using MAFFT v7.3. Multiple separate alignments were generated for large families such as GT2 and GT8 to capture the diversity within these families. These alignments made up the seed profiles for the GT-A families. These seed profiles were multiply aligned using the mapgaps scheme that allows alignment of diverse and large number of sequences using consensus sequence alignments and distinguishing aligned positions from insert segments. To guide the alignment of these profiles, a structure-based alignment of all available GT-A PDB sequences was built using Expresso of the Tcoffee suite (4) and MAFFT. Alignments for families with no representative crystal structures were guided using secondary structure predictions performed using PCI-SS (5). Finally, the alignment of secondary structures and conserved motifs were manually examined and corrected, where necessary.

During the generation of these profile alignments, we found several families that were very divergent and lacked nearly all canonical GT-A motifs. Since the features of GT-As are already so diverse, these divergent families such as family GT29 and GT42 sialyltransferases proved difficult to get a reasonable alignment with other families. Structural alignments also failed to properly align these families with other GT-A fold families. Thus, the following families were not included in the profile alignments: GT29, GT42 and GT46. These families are noted as atypical GT-A fold to distinguish them from other typical GT-A families that are part of this analysis. These families could represent atypical outliers from the GT-A fold which will be the subject of a separate analysis. This resulted in the inclusion of the 34 CAZy GT-A fold families in our profile alignments (Table S1).

#### Using the profile alignment to mine for and align putative GT-A fold sequences

Once the GT-A profiles were built, they were used in a sequence similarity search using mapgaps to identify and align more than 600,000 GT-A domain sequences from the NCBI non redundant sequence database. Sequences that had only short aligned segments (<40% aligned positions) were considered as fragmentary sequence hits and removed. Sequences with gaps at the canonical DxD motif positions were also removed from this alignment to exclude false hits and low confidence sequences.

#### Defining the GT-A common core

The profile alignments were used to describe the bounds for the core GT-A domain starting from the first beta sheet of the Rossmann fold domain to a C-terminal helix with family specific motifs. These bounds spanned the well-aligned conserved regions across all GT-A families. Within this common core, we identified regions that shared no overlap in structures and had the least sequence similarity across families to pinpoint the HVs. The conserved motifs and well aligned regions served as anchors to precisely define the bounds of these HVs. During our search, we found several classes of GT-A sequences that were multidomain and included either tandem GT domains (like the GT8 and GT49 LARGE domains) or other accessory domains (like the GT27 and lectin domains). We used our bound definitions above to separate out only the catalytic GT-A domains for such sequences. Multiple GT-A domains were separated and placed in their respective families. Such domains are referred to in the text as being either the N- or C-terminal domain based on their placement in the sequence.

#### Subset selection for Bayesian statistical analysis

To identify significantly conserved aligned positions across all GT-A fold families and identify positions conserved at a family specific level, we used the Bayesian procedure omcBPPS. Since Bayesian methods generally are very computationally intensive, running them on large datasets is not computationally feasible. Thus, we generated a representative subset of sequences that captured all the GT-A families and their taxonomic diversity. First, only single sequence was retained for sequences with more than 70% similarity. Then, larger CAZy families such as GT2 and GT8 were further trimmed to 50% similarity. Sequences from overrepresented taxonomic groups were further removed to reduce the dataset to a representative set of ~24,650 sequences.

#### Phylogenetic analysis

*Selection of sequences for phylogenetic analysis:*

Firstly, all identified GT-A sequences were selected form 5 model organisms: *H. sapiens*, *C. elegans*, *D. melanogaster*, *A. thaliana* and *S.cerevisiae.* For each of the 99 pattern-based groups, the highest scoring sequence for each identified taxonomic lineage (Archaea, Bacteria, Protista, Fungi, Metazoa, Plants, Virus) within these groups were selected next. For some GT-A families that did not have a representative pattern-based group, sequences were selected from their seed profiles. This resulted in a total of 993 sequences. These selected sequences are listed in Dataset S2 and their trimmed alignment used for phylogenetic inference is provided in Dataset S3.

*Details of the phylogenetic inference:*

IQTree v1.6.1 was used for phylogenetic inference with the following options: -nt AUTO -st AA -m MFP+MERGE -alrt 1000 -bb 1000 -wbt -nm 1000 -bnni. This implements ModelFinder (6) to select the best fit model based on Bayesian Information Criterion (BIC) and performs UltraFast Bootstrap (7) and the *SH-aLRT* (8) test with 1000 replicates to generate branch support. The -bnni option optimizes the bootstrap trees using a hill climbing nearest neighbor interchange

(NNI) search and reduces the risk of overestimating branch supports due to severe model violations.

*Generation of the condensed GT-A family phylogenetic tree (in Fig. 2) from the full GT-A tree (in Fig S4):*
The original phylogenetic inference was done using 993 sequences resulting in a large tree with 993 tips. Since visualization and interpretation of such a large tree is difficult, we condensed this large tree as follows: The deepest node that included all the sequences from the same family was collapsed such that the new tip generated at this node represents that GT-A family. For the GT-A families that did not form a monophyletic clade, the clade that included the most sequences from that family was used to represent the family tip. Sequences that did not fall within the tip clade were omitted during this collapse for clarity. While branch lengths were drawn to approximate the original distances, they were not drawn to scale. The full tree has been provided both as an image in Fig S4 and as a Newick file in Dataset S4 for a detailed overview of relationships at the sequence level. However, since most discussion in the paper are based on the GT-A family level relationship, Fig. 2 serves as the best visual representation of the tree for interpretability.

#### Orthogonal support for the phylogenetic tree

To test the robustness of the phylogenetic tree, its topology was compared to trees generated using orthogonal methods. Methods such as Hidden Markov Model (HMM) distances and structural similarities have been used in previous studies to estimate the phylogenetic relationships of subsets of GT-A families (9, 10). Here, we use a direct comparison of these methods to identify clades that show consistent hierarchical placement. The HMM-distance based phylogenetic tree was built using pHMM-Tree. Briefly, hmm profiles were built for each of the 53 sub-families identified in our analyses. Pairwise distances between these profiles were calculated and the resulting distance matrix was used to build a neighbor joining tree. All trees were visualized using the interactive Tree of Life (iTOL) online tool (11). Pairwise root mean square distance (RMSD) were calculated for 50 unique representative GT-A structures using the cealign algorithm in PyMol v2.0.6 (12) to build a distance matrix. Only the defined GT-A catalytic domain spanning the 231 aligned positions along with insertions were used for the RMSD calculations. This RMSD matrix was then used for clustering using the “ward” method in python which resulted in a structural distance based hierarchical clustering of the pdb structures. The hierarchical topology obtained from the HMM distance-based method and the RMSD distance based clustering were then compared to the tree topology in Fig. 2. This comparison is summarized in Fig S7. Connections that do not overlap connect consistently placed families, whereas families that are connected by connectors that overlap have been placed into different clusters by these different methods.

#### Machine learning analysis

*Gathering the training and validation dataset:*
For collecting the training set, we mined the “characterized” tab of the CAZy GT database as the primary source. Additionally, we also collected GT-A sequences from the UniProt database that

had known donors. Donor sugars for these characterized enzymes were identified based on their assigned names. Using this method, we curated a total of 713 GT-A domain sequences with known donor sugars that were then used as the training dataset for training the machine learning (ML) models. In addition, we also curated 64 GT-A sequences with known donor sugars for 5 model organisms (*H. sapiens*, *C. elegans*, *D. melanogaster*, *A. thaliana* and *S.cerevisiae*). These sequences were not used to train the ML model but set aside to be used as validation dataset to test the performance of the model.

Since the performance of ML methods relies heavily on the size of the training dataset, we had to merge 7 donor types (Ara, Fuc, GalF, GlcA, ManNAc, Rham, and Xyl) into a single category of "Others" as each of them had fewer than 50 sequences in the training dataset. This led us to build a 6-class classification model for predicting 6 major donor types: Glc, GlcNAc, Gal, GalNAc, Man, and "Others" with each class having more than 70 sequences in the training dataset. We then generated a multiple sequence alignment for the 713 training sequences by aligning them to our GT-A profiles using the mapgaps scheme described above. Out of the 231 aligned positions in this alignment, 147 positions were retained to remove highly gapped columns (removed positions with >15% gaps). For each of these 147 positions, we generated a set of five features using AAindex (13) to describe the property of the amino acid at these positions: charge, polarity, hydrophobicity, average accessible surface area, and side chain volume.

*Feature selection to build ML model:*

In order to select informative features to predict GT-A donor type from the 735 features (5 amino acid features for each of the 147 aligned positions), we implemented correlation-based feature selection (CFS) (14) with 5-fold CV by using WEKA version 3.8.3 (15) under default settings. Features selected by at least one iteration during the 5-fold CV were chosen. This led to the selection of 239 features for building multiple multiclass classification models. To further extract the most contributing features, we implemented CFS, information gain, and ReliefF by using WEKA with default settings. Each feature selection method with 10-fold CV was repeated five times and the results were aggregated. Among the most informative 50 features from each method, 24 of them were from at least two methods. To minimize Akaike information criterion (AIC) (16), the number of features was further narrowed down by using stepwise selection implemented by JMP® [JMP®, Version 14.1.0. SAS Institute Inc., Cary, NC, 1989-2019.]. As a result, 15 features were selected by multiple feature selection methods and determined to be the most significant contributing features.

*ML model training:*

We first trained random forest models by using an R package "randomForest" (17) with limited number of trees (ntree = 300) and limited maximum number of terminal nodes (maxnodes = 100) to avoid unrestricted tree expansion and potential overfitting. Two separate models were trained where the first one was trained with the larger set of 239 features and the second model was provided only 25 features coming from the donor binding residues. The importance of each feature used in the first random forest model was measured by the mean decrease in the Gini index (18). To compare the performance of this model, we also trained SVM, multilayer perceptron, Bayesian network, logistic regression, naive Bayes classifier, and J48 models by using WEKA with 10-fold CV under default settings. 10-fold CV evaluates the ML models by iteratively training on 90% of the data selected at random and testing the prediction on the unseen 10% of the data. This is repeated 10 times and the results on the testing dataset are summarized into an accuracy measure. The random forest model trained with 239 features had the highest accuracy and overall performance and thus was selected as the model of choice for predicting donor sugar substrates for GT-A enzymes.

*Evaluating the confidence of predictions:*

The confidence in prediction can be assessed by the probability scores assigned for each of the six prediction classes. The class with the highest probability represents the predicted donor sugar. As such, larger differences in probability between the first and second predicted class result in more reliable predictions. To interpret this difference in score easily, we derive a three-category confidence level. If the probability for the first class is more than double the probability of the second predicted class, then it is considered a high confidence prediction. If the difference is less than double but the probability of the first predicted class is more than double the probability by random chance, it is considered a moderate confidence prediction. If it is neither, then it is a low confidence prediction (Dataset S6).

#### Structural alignment of Rossmann fold proteins

Select representative set of structures were collected from all Rossmann-fold containing protein domains using the SCOP database (19). mTM-align (20) was used to align these structures with a subset of GT-A structures (Fig S1).

#### Construction of the sequence similarity network

A sequence similarity network was generated to evaluate the general predictability of a donor substrate based on sequence homology alone. Using an Edge-Weighted Spring Embedded Layout (21) in Cytoscape (22) with the same sequence dataset as the ML data, we produced a series of homologous networks constrained by an E-value cut-off of 0.05. Many families clustered on their own, and other families, such as GT31, GT8, and GT2 would be homologous enough to other families to cluster together. Within and between families, we noticed that sequences similar in identity, represented by nodes with short edges, could still select for vastly different donors. Certain families were highly conserved for their respective specific donor, while others varied significantly even amongst close homologs. This inconsistency made manual prediction via sequence homology an inappropriate method to perform donor prediction.

#### Selection of ancient archaeal and bacterial GT-A domain sequences in Dataset S7

The ancient archaeal and bacterial GT-A domain sequences listed in Dataset S7 represent sequences with the minimal GT-A domains (no long inserts, no additional domains) in prokaryotic organisms. These sequences could represent the most ancient progenitors of the GT-A fold families. These were selected based on closest homology to a consensus sequence derived from the GT-A profile alignments. First, a BLAST (23) search was conducted with the consensus sequence on then NCBI nr database. The 500 hits with an e-value better than 1e-10 were selected. These hits were then aligned to the GT-A profiles using mapgaps. Using this alignment, these sequences were further filtered to include hits that had 1) more than 140 aligned positions to remove fragmentary hits, 2) less than 20 insert positions throughout the GT-A domain alignment, 3) no more than 100 amino acids in the N-terminal region and 4) no more than 200 amino acids in the C-terminal region.

### Supplementary figures

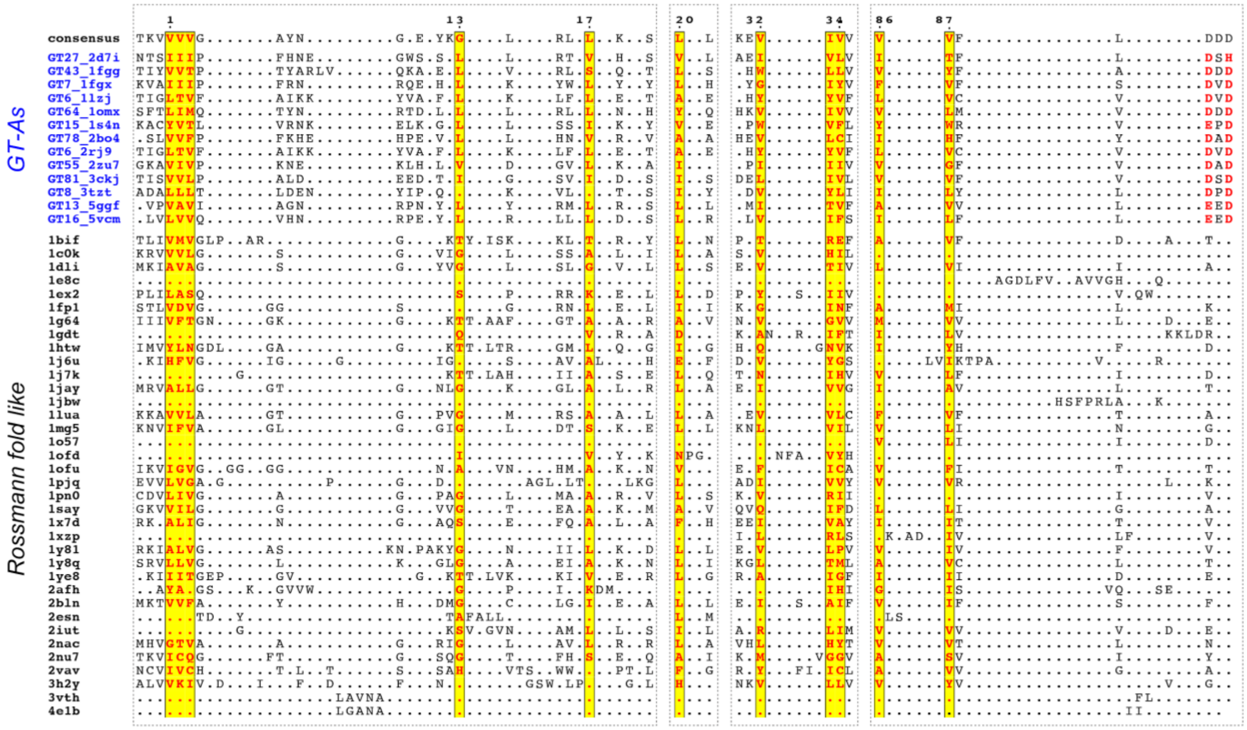

**Fig. S1**: Structure based sequence alignment showing the hydrophobic residue positions present across a collection of Rossmann fold like enzymes (highlighted in yellow blocks). Aligned positions are indicated at the top that correspond to aligned positions in Fig. 1D. The alignment extends until the DxD motif. Other regions were unaligned due to very low homology.

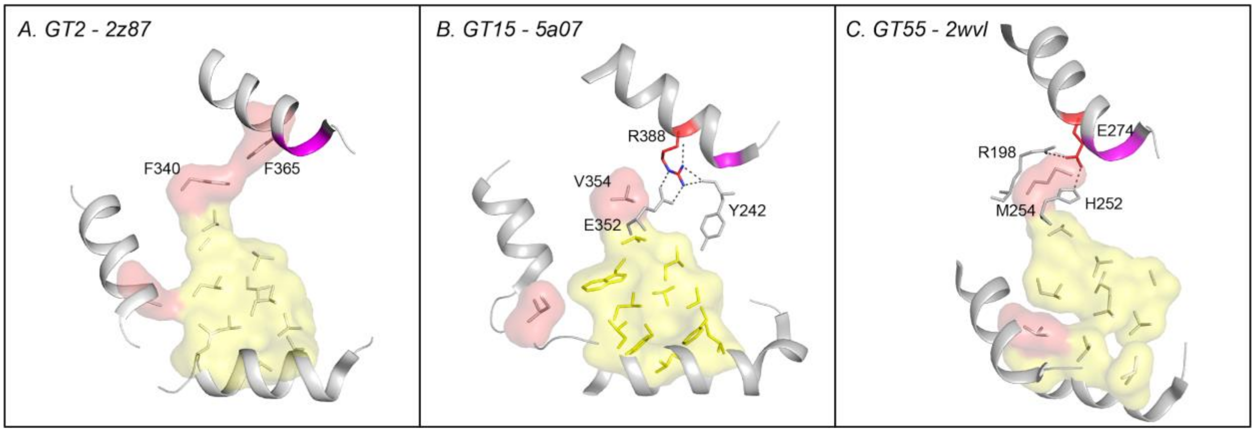


**Fig. S2**: Changes in the extended hydrophobic core residues in selected retaining families. A) The conserved hydrophobic core in the prototypic GT (2z87). B and C) Hydrophobic residue in the core is substituted by an Arginine and a Glutamate in GT15 and GT55 respectively. The charged residue replacing the hydrophobic residue of the core is highlighted in red sticks. The xED motif is shown in magenta.

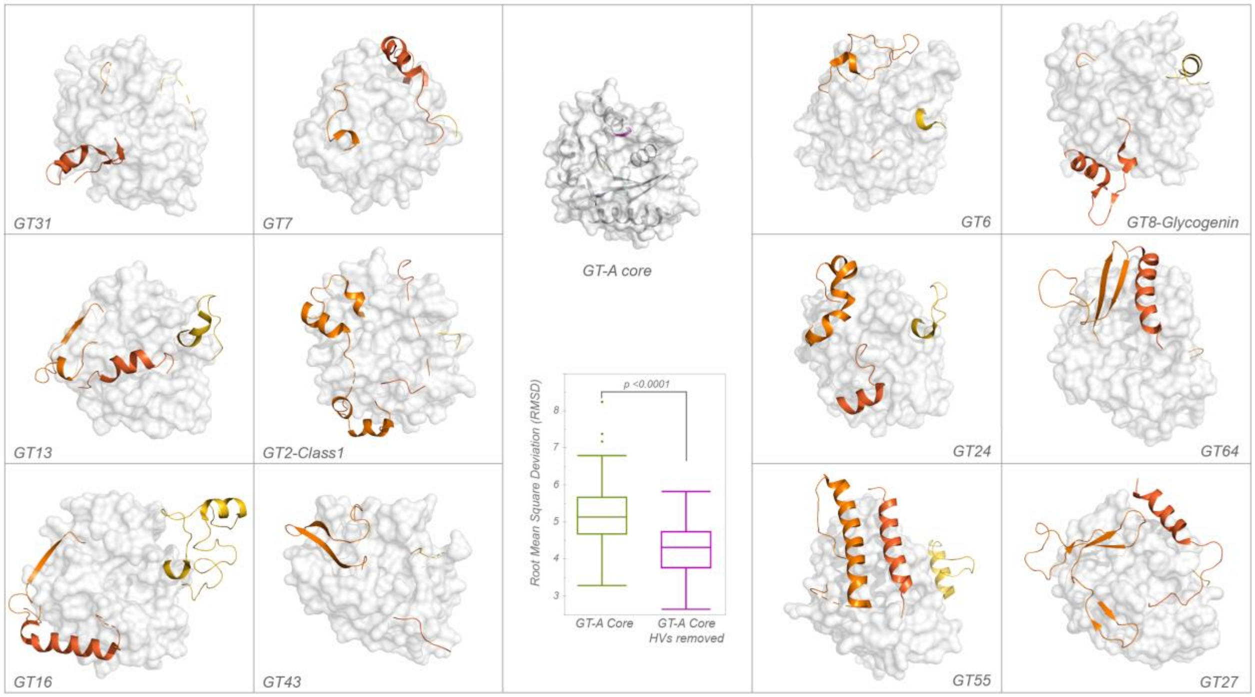


**Fig. S3**: Comparison of structures for HV regions across GT-A families. The GT-A common core is shown in surface in the middle. HVs are shown in shades of orange (HV1: light orange, HV2: dark orange, HV3: orange red). Root Mean Square Deviation (RMSD) was calculated by aligning the core GT-A domains of representative structures with and without the HVs. A significant reduction in the RMSD values was observed after removing HVs that is shown in the box plot in the center. *p-value <0.0001, t-test

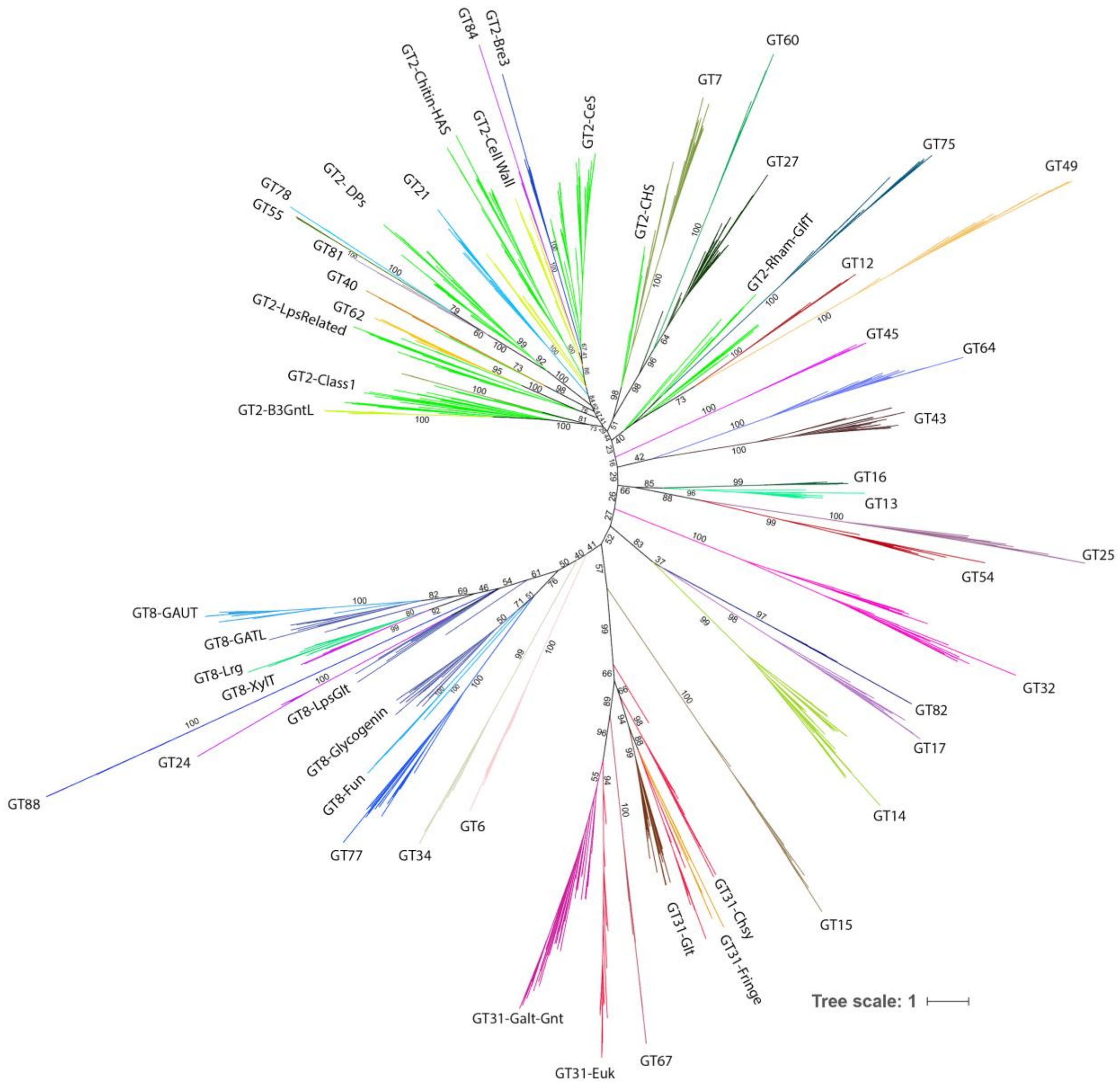


**Fig S4**: Complete phylogenetic tree of 993 representative GT-A sequences (Dataset S2). Clades are colored for each of the 53 GT-A families and labeled. Values at nodes indicate bootstrap support with 1000 replicates. Values for all major nodes are indicated. This tree is also provided in Newick text format in Dataset S4.

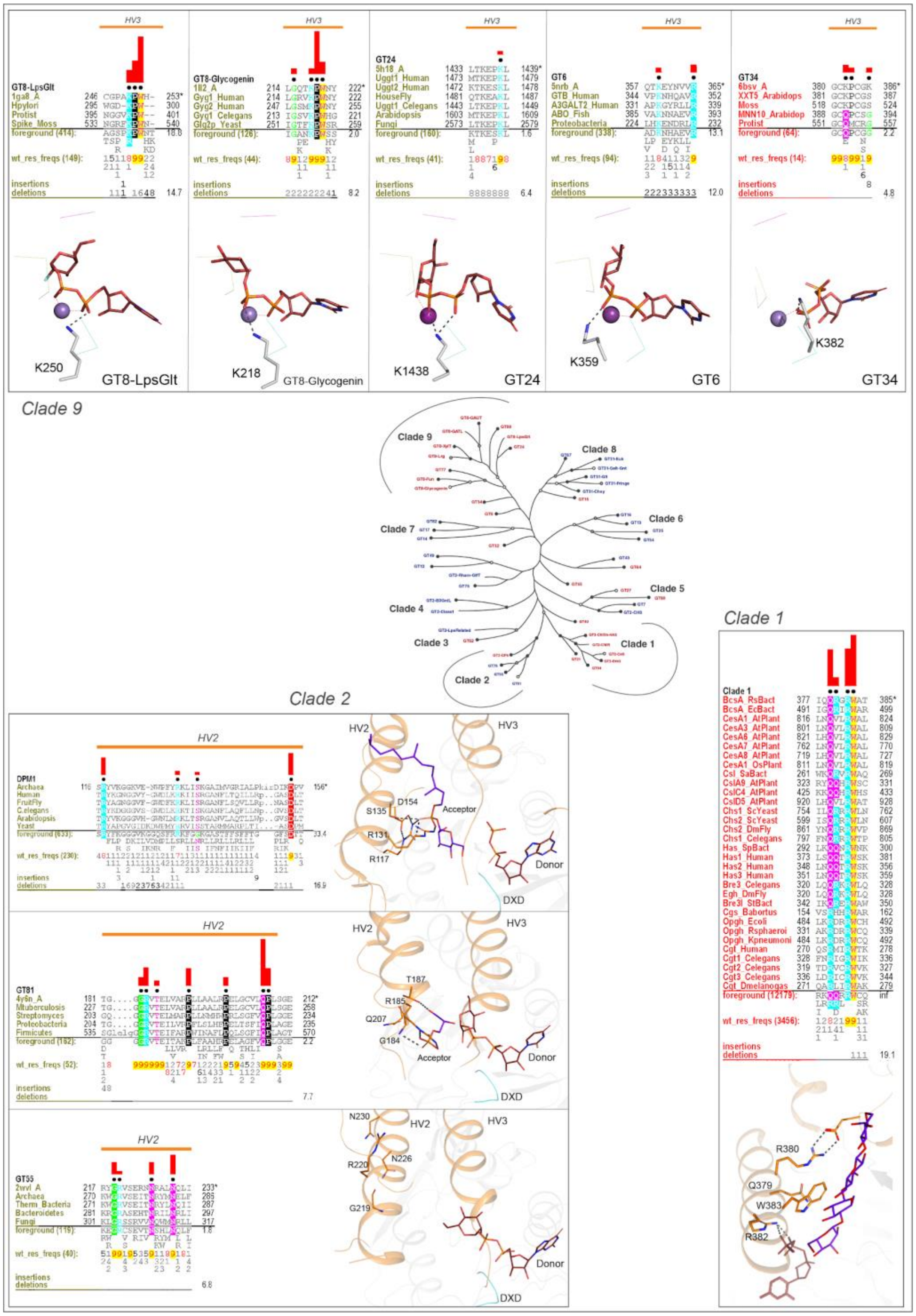


**Fig S5**: Clade specific conserved features in the HVs. The conserved mode of donor binding in clade 9, conserved mode of acceptor binding in clade 2 and the conserved QXXRW motif in clade 1 are illustrated. HVs are shown in orange. Metal ions are shown as spheres. Red bars above the alignment indicate the extent of significance of conservation of residue in the column (Higher is more significantly conserved). Below every position in the alignment, numbers indicate the extent of conservation of residues at the position.

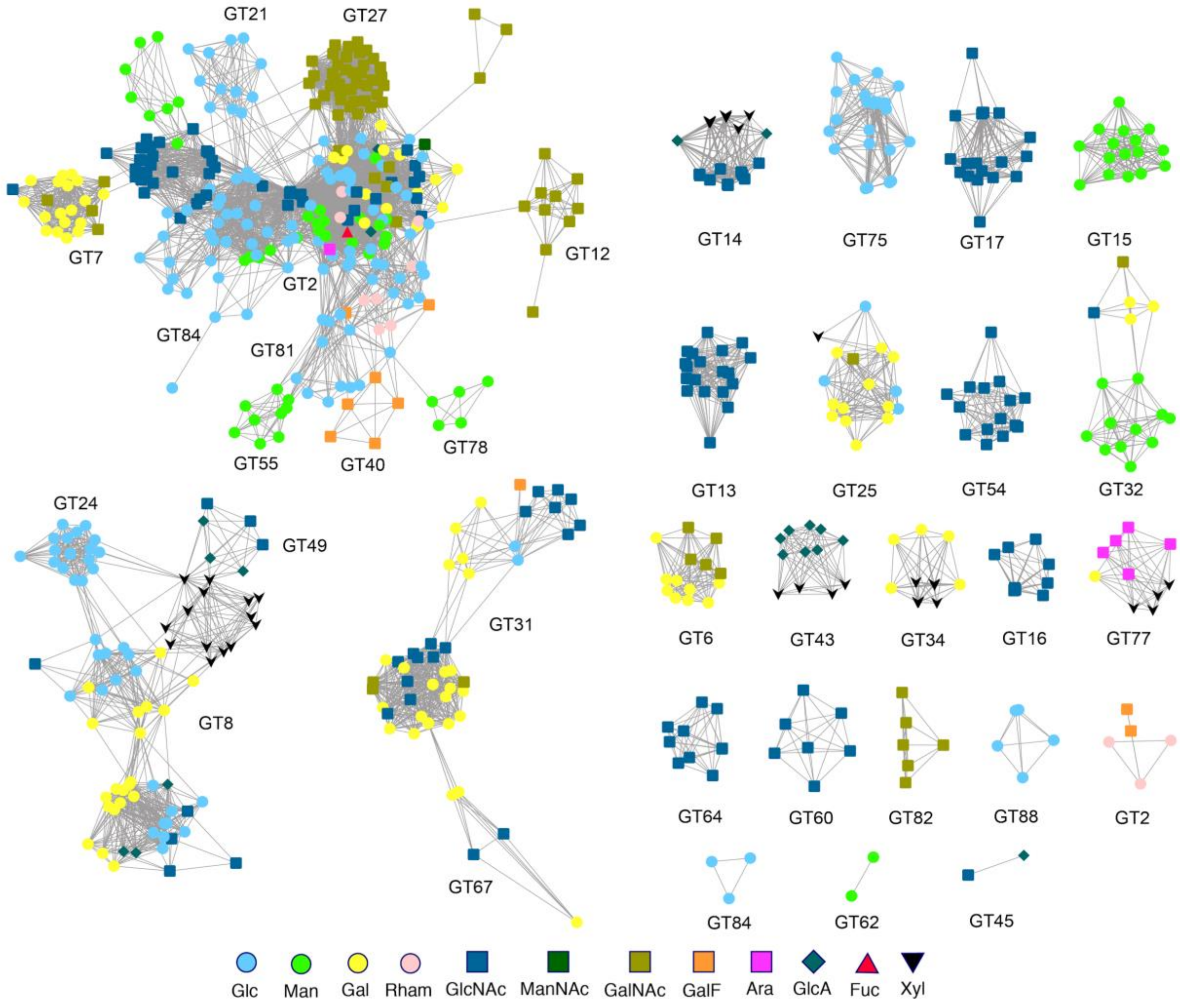


**Fig S6**: Sequence homology-based network of all the experimentally characterized sequences form the GT-A fold families (SI Methods). Nodes represent the sequences that were annotated as characterized and collected from the CAZy database to be used in the training dataset for ML. The color and shape of the nodes indicate the donor specificity for that sequence. An edge between two nodes indicate that the sequences are homologous with an e-value better than 1e-5. Smaller edge distance indicates a higher similarity between nodes. An edge-weighted spring embedded layout from Cytoscape was implemented to minimize edge crossings and enhance visual interpretability. At multiple locations in the network, closely related sequences differ in donor specificity, rendering prediction through similarity alone difficult.

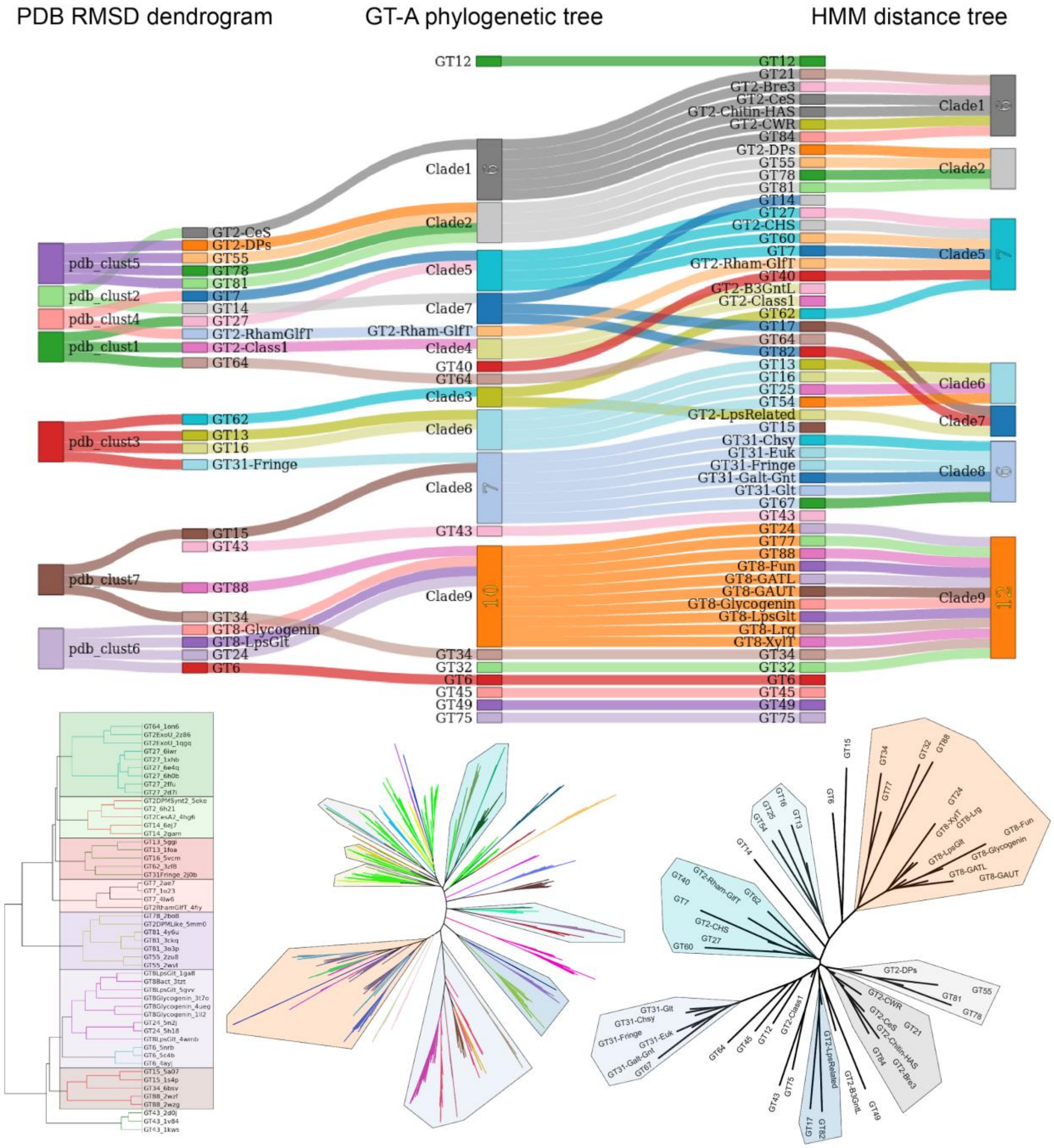


**Fig S7**: Sankey diagram comparing topologies of phylogenetic tree with pdb and hmm based clustering of GT-A families. Each column highlights clusters of GT-A families obtained through different methods (from left to right: PDB structural alignment clustering, GT-A phylogeny and hmm-distance based tree). Corresponding GT-A families within clusters are connected through colored links. Non overlapping links indicate an agreement in the placement of families across methods. Full clusters and trees are shown below the columns.

#### Supplementary tables

**Table S1: The CAZy GT-A families included in the analysis.** The "Alignment source" column includes the CDD alignment identifiers used to build the seed profiles for a given GT-A family. If a suitable CDD profile was not available, the seed profiles were built by manually selecting and aligning representative sequences.

| GTA Fam | Alignment source |
|---|---|
| GT2 | cd02511, cd02522, cd02525, cd02526, cd04184, cd04185, cd04187, cd04188, cd04190, cd04192, cd04195, cd04196, cd06420, cd06421, cd06427, cd06433, cd06434, cd06435, cd06437, cd06438, cd06439, cd06442, cd06913 and alignment of closest homologs of *C. elegans* bre3 |
| GT6 | cd02515 |
| GT7 | cd00899 |
| GT8 | cd02537, cd04194, cd06429, cd06430, cd06431, cd06914 and pfam01501 |
| GT12 | Alignment of selected CAZy sequences |
| GT13 | cd02514 |
| GT14 | Alignment of selected CAZy sequences |
| GT15 | pfam01793 |
| GT16 | pfam05060 |
| GT17 | Alignment of selected CAZy sequences |
| GT21 | cd02520 |
| GT24 | cd06432 |
| GT25 | cog3306 |
| GT27 | cd02510 |
| GT31 | pfam01762, pfam02434, pfam04646 and alignment of CAZy sequences |
| GT32 | Alignment of selected CAZy sequences |
| GT34 | pln03182 |
| GT40 | Alignment of selected CAZy sequences |
| GT43 | cd00218,pfam03360 and alignment of CAZy sequences |
| GT45 | Alignment of selected CAZy sequences |
| GT49 | pfam13896 |
| GT54 | pfam04666 |
| GT55 | pfam09488 |
| GT60 | pfam11397 |
| GT62 | Alignment of selected CAZy sequences |
| GT64 | pfam09258 |
| GT67 | Alignment of selected CAZy sequences |
| GT75 | Alignment of selected CAZy sequences |
| GT77 | Alignment of selected CAZy sequences |
| GT78 | Alignment of selected CAZy sequences |
| GT81 | prk13915 |
| GT82 | pfam06306 |
| GT84 | cd04191 |
| GT88 | Alignment of selected CAZy sequences |

**Table S2: List of GT-A fold families and subfamilies.** For each of these families, the groups obtained by the pattern based classification are provided in the "GT-A pattern based group" column. Taxonomic distribution and a short description of these groups are also provided.

| CAZy family | GT-A family | GT-A pattern based group | Description | Number of sequences in NCBInr | Taxonomic distribution | | | | | | | |
|---|---|---|---|---|---|---|---|---|---|---|---|---|
| | | | | | Archaea | Bacteria | Protista | Fungi | Metazoa | Viridiplantae | Viruses | Unknown |
| GT2 | GT2-B3GntL | GT2-B3GntL-1 | beta-1,3-N-acetylglucosaminyltransferase-like sequences from GT2 | 683 | - | - | 51 | - | 621 | 7 | - | 4 |
| | GT2-Bre3 | GT2-Bre3-1 | Close homologs of *C. elegans* bre3 and Drosophila egh GT2 sequences | 696 | 184 | 106 | 11 | 125 | 264 | 1 | - | 5 |
| | GT2-CWR | GT2-CWR_Csl-1 | 1st subgroup of GT2-CWR sub-family that includes bacterial Cellulose synthase like sequences | 2229 | - | 2223 | - | - | 2 | - | - | 4 |
| | | GT2-CWR_Csl-2 | 2nd subgroup of GT2-CWR sub-family that includes bacterial and archaeal Cellulose synthase like sequences | 3386 | 21 | 3363 | - | 2 | - | - | - | - |
| | GT2-CeS | GT2-CeS-1 | Cellulose synthases, Cellulose synthase like and related sequences | 4142 | 102 | 1768 | 9 | - | 6 | 2255 | - | 2 |
| | | GT2-CeS_putative | Mostly bacterial putative cellulose synthesis related sequences | 2112 | 1 | 2109 | - | - | - | - | - | 2 |
| | GT2-Chitin-HAS | GT2-Chitin-HAS-1 | Chitin and hyaluronan synthases of the GT2 family | 15602 | 99 | 4813 | 231 | 8357 | 2005 | 38 | 42 | 17 |
| | GT2-Class1 | GT2-Class1-1 | Large subgroup of GT2 sequences most with unknown functions | 17387 | 143 | 17176 | 36 | - | 2 | 2 | 2 | 26 |
| | | GT2-Class1-2 | Large subgroup of GT2 sequences most with unknown functions | 7496 | 15 | 7455 | 1 | - | 2 | 1 | - | 22 |
| | GT2-DPs | GT2-DPs_Bact | Mostly bacterial sequences that transfer sugar to Dolichol Phosphate acceptors (DPG/DPM) | 34163 | 106 | 33968 | 2 | 4 | 12 | 6 | 6 | 59 |
| | | GT2-DPs_Mix | Mix of Dolichol phosphate glucose and mannose (DPG/DPM) glycosyltransferases | 66769 | 3901 | 58998 | 300 | 1531 | 1376 | 479 | 2 | 182 |
| | GT2-LpsRelated | GT2-LpsRelated-1 | Prokaryotic GT2 sequences predicted to be involved in cell wall biosynthesis | 25139 | 69 | 24998 | 3 | - | 4 | 10 | 24 | 31 |
| GT6 | GT6 | GT6-Mix | homologs of GBGT1 GalNAc transferases and bacterial GT6 sequences | 1067 | 1 | 211 | 4 | 3 | 834 | 2 | 10 | 2 |
| | | GT6-A3GALT | Homologs of Alpha 3 Galactosyltransferases (A3GALT) | 704 | - | - | - | - | 703 | - | - | 1 |
| | | GT6-ABO | Blood ABO transferases | 839 | - | - | - | - | 835 | - | - | 4 |
| | | GT6-GLT6D1 | GLT6D1 related sequences | 226 | - | - | - | - | 224 | - | - | 2 |
| GT7 | GT7 | GT7-1 | All GT7 sequences | 4372 | 5 | 62 | 38 | 1 | 4238 | 7 | 4 | 17 |
| GT8 | GT8-Fun | GT8-Fun-1 | Fungi and some bacterial GT8 sequences | 738 | - | - | 28 | 701 | - | 9 | - | - |
| | GT8-GATL | GT8-GATL-1 | Mostly plant alpha-galacturonosyltransferase like sequences | 144 | - | - | 4 | - | 6 | 134 | - | - |
| | GT8-GAUT | GT8-GAUT-1 | Mostly chordate and plant alpha-galacturonosyltransferase homologs | 906 | - | - | 5 | 1 | 96 | 804 | - | - |
| | | GT8-GAUT-2 | Alpha-galacturonosyltransferase homologs from multiple taxa | 1714 | - | - | 8 | - | 106 | 1600 | - | - |

| | | | | | | | | | | | | |
|---|---|---|---|---|---|---|---|---|---|---|---|---|
| | | GT8-GAUT-3 | Plant alpha-galacturonosyltransferase homologs | 503 | - | - | - | - | - | 503 | - | - |
| | | GT8-GAUT-4 | alpha-galacturonosyltransferase homologs from multiple taxa | 938 | - | - | - | - | 22 | 916 | - | - |
| | GT8-Glycogenin | GT8-Glycogenin-1 | Non-metazoan Glycogenin related sequences | 1143 | 1 | 194 | 39 | 895 | 1 | 10 | 2 | 1 |
| | | GT8-Glycogenin-2 | Glycogenin related sequences from all taxa | 2937 | - | 51 | 34 | 1022 | 1778 | 17 | 21 | 14 |
| | GT8-Lrg | GT8-Lrg-1 | Eukaryotic GT8 Large domain xylosyltransferase homologs | 1204 | - | 1 | 13 | 19 | 1156 | - | - | 15 |
| | GT8-XylT | GT8-XylT-1 | Eukaryotic GT8 xylosyltransferases | 1252 | - | - | 4 | 27 | 1212 | 4 | - | 5 |
| GT12 | GT12 | GT12-1 | All GT12 sequences | 1018 | 4 | 65 | 12 | 2 | 933 | - | - | 2 |
| GT13 | GT13 | GT13-POMGNT1 | POMGNT1 homologs | 1272 | - | 225 | 85 | 2 | 663 | 295 | - | 2 |
| | | GT13-MGAT1 | MGAT1 homologs | 727 | - | - | 2 | 1 | 719 | - | - | 5 |
| GT14 | GT14 | GT14-1 | All GT14 sequences | 8206 | 1 | 2790 | 77 | 11 | 3061 | 2229 | 10 | 27 |
| GT15 | GT15 | GT15-1 | All GT15 sequences | 3547 | - | 12 | 26 | 3469 | 16 | 23 | 1 | - |
| GT16 | GT16 | GT16-1 | Protist, Plants, Nematode and other Metazoan MGAT2 homologs | 499 | - | - | 14 | - | 275 | 210 | - | - |
| | | GT16-2 | Arthropod and Chordate MGAT2 homologs | 648 | - | - | - | - | 645 | - | - | 3 |
| GT17 | GT17 | GT17-1 | All GT17 sequences | 2672 | 5 | 750 | 40 | 501 | 540 | 823 | 9 | 4 |
| GT21 | GT21 | GT21-1 | Mostly Prokaryotic members of the GT21 family | 2958 | 72 | 2843 | 3 | 1 | 3 | 34 | - | 2 |
| | | GT21-2 | Fungi nearly all Ascomycota GT21 members | 458 | - | - | - | 455 | - | 3 | - | - |
| | | GT21-3 | Fungi GT21 sequences | 324 | - | - | - | 324 | - | - | - | - |
| | | GT21-4 | Proteobacteria + Metazoa GT21 sequences | 1331 | - | 737 | 2 | - | 587 | - | - | 5 |
| GT24 | GT24 | GT24-1 | All GT24 sequences | 2756 | - | - | 139 | 935 | 1397 | 279 | - | 6 |
| GT25 | GT25 | - | All GT25 sequences; No pattern based group | 12296 | - | 10425 | 94 | 198 | 1450 | 21 | 96 | 12 |
| GT27 | GT27 | GT27-1 | All GT27 sequences | 11047 | 2 | 186 | 193 | - | 10618 | 1 | - | 47 |
| GT31 | GT31-Chsy | GT31-Chsy-1 | CHSY1 and CSS3 homologs | 893 | - | - | 7 | 11 | 870 | 2 | - | 3 |
| | | GT31-Chsy-2 | Metazoan CHPF homologs | 898 | - | - | 3 | - | 890 | - | 2 | 3 |
| | GT31-Euk | GT31-Euk-1 | Mostly plant GT31 sequences with unknown functions | 721 | - | 1 | - | - | - | 720 | - | - |
| | | GT31-Euk-2 | Mostly plant GT31 sequences; Includes Avr9 elicitor from *A. thaliana* | 1979 | - | - | - | - | - | 1979 | - | - |
| | | GT31-Mix-1 | Mix of GT31 sequences from all taxonomic groups; Includes Sqv-2 from *C. elegans* | 538 | - | 6 | 14 | 18 | 492 | 7 | - | 1 |
| | GT31-Fringe | GT31-Fringe-1 | GT31 Fringe homologs | 1438 | - | - | 1 | 1 | 1411 | 18 | - | 7 |
| | GT31-Glt | GT31-Glt_B3GLCT | Eukaryotic B3GLCT homologs | 981 | - | - | 8 | 5 | 777 | 188 | - | 3 |
| | | GT31-Glt_C1GALT | Includes C1GALT1 and its close homologs | 2153 | - | 33 | 169 | 52 | 1867 | 22 | - | 10 |

| | | | | | | | | | | | | |
|---|---|---|---|---|---|---|---|---|---|---|---|---|
| GT32 | GT32 | GT32-1 | alpha 1,4-galactosyltranseferases from *A. thaliana* and plant GT32 sequences | 585 | - | 6 | 1 | 17 | - | 561 | - | - |
| | | GT32-2 | A4GALT1 and 2 from *D. melanogaster* and other unknown GT32 sequences | 685 | 1 | 27 | 6 | 84 | 402 | 165 | - | - |
| | | GT32-3 | A4GNT and A4GALT from human and mostly chordates homologs | 746 | - | - | - | 6 | 737 | - | - | 3 |
| | | GT32-4 | Mostly bacteria, protozoa and fungi GT32 sequences | 4738 | - | 1547 | 129 | 2839 | 109 | 62 | 51 | 1 |
| | | GT32-5 | Mostly bacteria GT32 sequences | 2393 | - | 2365 | - | 6 | 6 | - | 8 | 8 |
| GT34 | GT34 | GT34-1 | All GT34 sequences | 1198 | - | 1 | 5 | 83 | 3 | 1106 | - | - |
| GT40 | GT40 | GT40-1 | All GT40 sequences | 167 | - | - | 167 | - | - | - | - | - |
| GT43 | GT43 | GT43-B3GAT3 | B3GAT3 homologs | 1865 | - | - | 20 | 30 | 836 | 974 | - | 5 |
| | | GT43-Mix-1 | Small subset of unknown GT43 sequences | 79 | - | - | - | - | 75 | 4 | - | - |
| | | GT43-B3GAT2 | Chordate homologs of B3GAT2 | 129 | - | - | - | - | 128 | - | - | 1 |
| | | GT43-Mix-2 | Lower metazoan and plant GT43 sequences | 125 | - | - | - | - | 122 | 3 | - | - |
| | | GT43-B3GAT1 | B3GAT1 homologs | 690 | - | - | - | - | 690 | - | - | - |
| | | GT43-Nematode | Nematode GT43 sequences | 73 | - | - | - | - | 73 | - | - | - |
| GT45 | GT45 | GT45-1 | All GT45 sequences | 410 | 6 | 395 | 6 | - | - | 1 | - | 2 |
| GT49 | GT49 | GT49-B4GAT1like | B4GAT1 homologs | 1353 | - | 1 | 183 | 50 | 1045 | 68 | - | 6 |
| | | GT49-Fungi | Fungi GT49 sequences of unknown function | 298 | - | - | - | 298 | - | - | - | - |
| | | GT49-LARGE | Homologs of human LARGE1 and 2 GT49 domains | 1111 | - | - | 5 | - | 1095 | 2 | - | 9 |
| GT54 | GT54 | GT54-MGAT4C | MGAT4C homologs | 1214 | - | 1 | 16 | 13 | 1176 | 5 | - | 3 |
| | | GT54-MGAT4AB | Closest homologs of MGAT4A and 4B | 1313 | - | - | - | - | 1310 | - | - | 3 |
| | | GT54-Chordate | Chordate GT54 sequences | 283 | - | - | 1 | - | 280 | - | - | 2 |
| GT55 | GT55 | GT55-1 | All GT55 sequences | 179 | 72 | 66 | - | 39 | - | - | - | 2 |
| GT60 | GT60 | GT60-1 | All GT60 sequences | 1125 | - | 616 | 469 | 3 | - | 26 | 11 | - |
| GT62 | GT62 | GT62-1 | Subgroup of GT62 sequences from all taxa | 383 | - | 156 | 57 | 145 | 17 | 7 | - | 1 |
| | | GT62-2 | Includes Mnn9p from *S. cerevesiae* and its close homologs | 566 | - | - | - | 562 | - | 4 | - | - |
| | | GT62-3 | Mostly fungal GT62 sequences | 1201 | - | - | - | 1193 | - | 8 | - | - |
| GT64 | GT64 | GT64-EXTLs | Exostatin like sequences | 1502 | - | 2 | 50 | 239 | 1042 | 162 | - | 7 |
| | | GT64-EXTs | Metazoan Exostatin sequences | 1637 | - | - | - | - | 1623 | - | - | 14 |
| | | GT64-Mix | Mix of GT64 sequence from diverse organisms, mostly plants | 599 | - | - | 20 | 3 | 1 | 575 | - | - |
| GT67 | GT67 | - | All GT67 sequences; Group with GT31 sequences in GT31-67-Mix | 56 | - | - | 56 | - | - | - | - | - |

| | | | | | | | | | | | | |
|---|---|---|---|---|---|---|---|---|---|---|---|---|
| GT75 | GT75 | GT75-1 | Prokaryote and plant GT75 sequences | 112 | 7 | 64 | 1 | - | - | 39 | - | 1 |
| | | GT75-2 | GT75 sequences from archaea, chlorophytes and plants | 1154 | 230 | 20 | - | - | 1 | 901 | - | 2 |
| | | GT75-3 | GT75 sequences from diverse taxonomic groups | 475 | - | 151 | 21 | 10 | 110 | 183 | - | - |
| GT77 | GT77 | GT77-1 | GT77 sequences from mix taxa | 1923 | 1 | 27 | 243 | 145 | 139 | 1366 | 2 | - |
| | | GT77-2 | Mostly plant GT77 sequences | 1402 | - | - | 1 | 1 | 2 | 1398 | - | - |
| GT78 | GT78 | - | All GT78 sequences; No pattern based group | 8 | - | - | 5 | - | - | 3 | - | - |
| GT81 | GT81 | GT81-1 | GT81 from Euryarchaeota and Cyanobacteria phyla | 472 | 251 | 217 | - | - | - | - | - | 4 |
| | | GT81-2 | GT81 sequences from archaea and bacteria | 3299 | 30 | 3258 | - | 1 | - | - | - | 10 |
| | | GT81-3 | Mostly proteobacteria GT81 sequences | 1035 | 1 | 1032 | - | - | - | - | - | 2 |
| GT82 | GT82 | GT82-1 | All GT82 sequences | 798 | - | 795 | 3 | - | - | - | - | - |
| GT84 | GT84 | GT84-1 | 1st subgroup with a mix of GT84 sequences | 7993 | 1 | 7927 | - | 49 | 8 | 2 | - | 6 |
| | | GT84-2 | 2nd subgroup with a mix of GT84 sequences | 738 | - | 736 | 1 | - | - | - | - | 1 |
| GT88 | GT88 | - | All GT88 sequences; No pattern based group | 184 | - | 184 | - | - | - | - | - | - |
| Mixed groups | | GT2-Mix-1 | Mix of sequences from multiple GT2 subfamilies, mostly prokaryotes | 39143 | 898 | 38072 | 12 | 29 | 22 | 22 | 2 | 86 |
| | | GT2-Mix-2 | Mix of mostly cellulose synthase like and CWR sequences. Also includes some GT84 and other GT2 sequences. | 40778 | 832 | 33292 | 203 | 994 | 23 | 5290 | 73 | 71 |
| | | GT2-Mix-3 | Mix of sequences from multiple GT2 subfamilies; mostly GT2-Class1 and GT2-RhamGlfT | 69424 | 1434 | 67547 | 22 | 42 | 214 | 16 | 1 | 148 |
| | GT2-CHS | GT2-Mix-4 | Mix of sequences mostly from the GT2-Class1 subfamily; Also contains sequences from the GT2-CHS and GT2-LpsRelated sub families | 117208 | 1620 | 115181 | 60 | 69 | 26 | 15 | 20 | 217 |
| | GT2-Rham-GlfT | GT2-Mix-5 | Majority are rhamnosyl and galactofuranosyl transferases from the GT2 family but also includes some sequences from GT2-Class1 sub family. | 23498 | 382 | 22962 | 1 | - | 48 | 2 | 71 | 32 |
| | GT31-Galt-Gnt | GT31-67_Mix | Mix of sequence homologs of B3GALTs, B3GNTs; and GT67 | 9546 | - | 21 | 579 | 29 | 7695 | 1190 | 2 | 30 |
| | | GT8-GAUT_GATL_Mix | Mix of GAUT and GATL like sequences | 1877 | - | - | 6 | 4 | 563 | 1304 | - | - |
| | | GT8-Mix-1 | Mix of GT8 sequences from GT8-Glycongenin, GT8-LpsGlt and GT8-GATL families | 4367 | 1 | 782 | 219 | 618 | 50 | 2630 | 66 | 1 |
| | GT8-LpsGlt | GT8-Mix-2 | GT8 sequences involved in lipopolysaccharide biosynthesis | 19773 | 26 | 19598 | 43 | 62 | 11 | 10 | 4 | 19 |
| | | GT8-Mix-3 | Metazoan GT8 sequences | 375 | - | - | 3 | - | 370 | - | - | 2 |
| | | GT2-DPs_81_Mix | Mix of GT2 dolichol phosphate mannose/glucose and GT81 sequences | 1543 | 94 | 1442 | - | - | - | 4 | - | 3 |

**Table S3: List of the top 100 of the 239 contributing features used in the random forest model ranked based on the Mean decrease in Gini Index.** For any given position, the 5 features of the amino acid at that position are VOL:Volume; POL: Polarity; ASA: Accessible Surface Area; CHA: Charge; HYD: Hydrophobicity. The "In 15 Most Contributing Features" indicates whether a feature was selected in the 15 most contributing features.

| Rank | Feature | Mean Decrease Gini | In 15 most Contributing Features |
|---|---|---|---|
| 1 | Pos176_VOL | 3.2262624 | Yes |
| 2 | Pos177_VOL | 3.1756798 | Yes |
| 3 | Pos92_VOL | 2.7699882 | Yes |
| 4 | Pos68_ASA | 2.6025505 | Yes |
| 5 | Pos165_VOL | 2.4503841 | - |
| 6 | Pos67_POL | 2.4325555 | - |
| 7 | Pos176_POL | 2.2984334 | - |
| 8 | Pos188_VOL | 2.2537245 | - |
| 9 | Pos192_POL | 2.1060804 | - |
| 10 | Pos67_ASA | 2.0168254 | Yes |
| 11 | Pos20_HYD | 2.0065955 | - |
| 12 | Pos182_ASA | 2.0015905 | - |
| 13 | Pos180_ASA | 1.9937479 | - |
| 14 | Pos69_VOL | 1.9770843 | - |
| 15 | Pos183_VOL | 1.8706995 | - |
| 16 | Pos24_VOL | 1.8449087 | - |
| 17 | Pos70_POL | 1.8443468 | - |
| 18 | Pos119_POL | 1.8106287 | - |
| 19 | Pos70_VOL | 1.7231085 | - |
| 20 | Pos165_HYD | 1.7092602 | - |
| 21 | Pos171_ASA | 1.7043347 | - |
| 22 | Pos68_POL | 1.6255744 | - |
| 23 | Pos177_ASA | 1.6069533 | - |
| 24 | Pos153_POL | 1.5926813 | - |
| 25 | Pos119_HYD | 1.5891478 | - |
| 26 | Pos154_HYD | 1.5841736 | - |
| 27 | Pos148_ASA | 1.5748568 | - |
| 28 | Pos181_VOL | 1.5053581 | - |
| 29 | Pos209_VOL | 1.4993641 | - |
| 30 | Pos48_POL | 1.4944802 | - |
| 31 | Pos31_VOL | 1.4819922 | - |
| 32 | Pos69_HYD | 1.4794809 | Yes |
| 33 | Pos183_POL | 1.4780106 | - |
| 34 | Pos95_VOL | 1.4297656 | - |
| 35 | Pos207_VOL | 1.4202223 | Yes |
| 36 | Pos66_ASA | 1.4158461 | - |
| 37 | Pos208_ASA | 1.3807237 | - |
| 38 | Pos148_VOL | 1.3782629 | - |
| 39 | Pos5_VOL | 1.361738 | - |
| 40 | Pos192_ASA | 1.3513478 | - |
| 41 | Pos119_ASA | 1.3459896 | - |
| 42 | Pos4_ASA | 1.3455248 | Yes |
| 43 | Pos71_ASA | 1.3343774 | Yes |
| 44 | Pos5_HYD | 1.329734 | - |
| 45 | Pos90_HYD | 1.3206017 | - |
| 46 | Pos77_POL | 1.3150915 | - |
| 47 | Pos150_HYD | 1.2936144 | Yes |
| 48 | Pos118_HYD | 1.2828461 | - |
| 49 | Pos171_CHA | 1.2745598 | - |
| 50 | Pos193_POL | 1.2571925 | - |
| 51 | Pos75_POL | 1.2562786 | - |
| 52 | Pos113_VOL | 1.2538753 | - |
| 53 | Pos184_HYD | 1.2439932 | Yes |
| 54 | Pos152_HYD | 1.240217 | - |
| 55 | Pos185_ASA | 1.2330839 | - |
| 56 | Pos16_VOL | 1.2296393 | - |
| 57 | Pos183_HYD | 1.2262149 | - |
| 58 | Pos96_VOL | 1.2101717 | - |
| 59 | Pos193_ASA | 1.2054209 | - |
| 60 | Pos169_POL | 1.1989623 | - |
| 61 | Pos71_VOL | 1.1979185 | - |
| 62 | Pos34_ASA | 1.1953853 | - |
| 63 | Pos65_POL | 1.1897809 | - |
| 64 | Pos34_POL | 1.1878609 | - |
| 65 | Pos69_POL | 1.1804113 | - |
| 66 | Pos195_VOL | 1.1777859 | - |
| 67 | Pos32_ASA | 1.1775224 | - |
| 68 | Pos67_VOL | 1.1677597 | - |
| 69 | Pos152_ASA | 1.1667168 | - |
| 70 | Pos36_VOL | 1.1660744 | - |
| 71 | Pos120_POL | 1.1643192 | - |
| 72 | Pos181_POL | 1.1619564 | - |
| 73 | Pos153_ASA | 1.1608432 | - |
| 74 | Pos174_VOL | 1.1555382 | - |
| 75 | Pos148_HYD | 1.1547405 | - |
| 76 | Pos75_VOL | 1.131398 | - |
| 77 | Pos88_VOL | 1.1308913 | - |
| 78 | Pos165_ASA | 1.108977 | - |
| 79 | Pos181_ASA | 1.1089336 | - |
| 80 | Pos179_ASA | 1.100562 | - |
| 81 | Pos70_ASA | 1.0932475 | - |
| 82 | Pos180_POL | 1.0915233 | - |
| 83 | Pos94_HYD | 1.0899666 | - |
| 84 | Pos184_VOL | 1.0877394 | - |
| 85 | Pos192_HYD | 1.0854422 | - |
| 86 | Pos115_HYD | 1.081659 | - |
| 87 | Pos102_POL | 1.0807846 | - |
| 88 | Pos88_ASA | 1.0807419 | - |
| 89 | Pos31_POL | 1.0661381 | - |
| 90 | Pos77_HYD | 1.0624357 | - |
| 91 | Pos72_POL | 1.0599151 | |
| 92 | Pos209_HYD | 1.0520118 | |
| 93 | Pos91_HYD | 1.0510114 | |
| 94 | Pos68_HYD | 1.0417099 | |
| 95 | Pos102_HYD | 1.038828 | |
| 96 | Pos24_POL | 1.0363453 | |
| 97 | Pos78_HYD | 1.0347368 | |
| 98 | Pos117_HYD | 1.0326644 | |
| 99 | Pos185_POL | 1.0224647 | |
| 100 | Pos58_ASA | 1.0213281 | |

#### SI Datasets

**Dataset S1:** Mapping of the 231 aligned positions to representative crystal structures available for all GT-A fold families. The "Position Description" column includes the labels for conserved motifs and hypervariable regions for the aligned positions. GT family, PDB IDs and reference to PubMed IDs are indicated at the header columns.

**Dataset S2:** The 993 representative GT-A domain sequences included in the phylogenetic analysis. The GT-A family and the pattern based classification group for each sequence is indicated in the "GT-A family" and the "GT-A pattern based group" columns. The domain start and end positions are indicated. Sequence for the domain region and the full length sequences are also provided. An alignment of these sequences are available in Dataset S3.

**Dataset S3:** The trimmed alignment of the 231 positions of the GT-A core used for phylogeny in FASTA format.

**Dataset S4:** The phylogenetic tree file for the 993 GT-A fold sequences in Newick format.

**Dataset S5:** List of the 713 training dataset sequences used for machine learning. The "Assigned Donor Class" column indicates one of the 6 classes the donor belongs to.

**Dataset S6:** The prediction results for donor prediction using the random forest ML model for GT-A sequences from five model organisms. The validation datasets (highlighted in blue rows) include GTs that have some experimental characterization but were not included in the characterized dataset. The validation set was used to compare the model predictions with the experimental results. The "Match Experimental" column indicates whether the prediction matched experimental results. The prediction set includes predictions for GTs of unknown functions. The "Confidence" column includes the confidence for prediction which was derived based on the probability for the 1st class and its difference with the probability for the 2nd class. Probabilities for all the 6 classes are provided in the "Classwise Probablity" columns.

**Dataset S7:** The ancient archeal and bacterial sequences that most closely resemble the GT-A core consensus. These sequences were collected by running a BLAST search against a single consensus sequence generated from the seed profiles of all GT-A families. The hits were further filtered to keep sequences that only had the minimal GT-A core (SI Methods).
